# The synapsin-dependent vesicle cluster is crucial for presynaptic plasticity at a glutamatergic synapse in male mice

**DOI:** 10.1101/2023.08.08.549335

**Authors:** Felicitas Bruentgens, Laura Moreno Velasquez, Alexander Stumpf, Daniel Parthier, Jörg Breustedt, Fabio Benfenati, Dragomir Milovanovic, Dietmar Schmitz, Marta Orlando

**Affiliations:** Charité – Universitätsmedizin Berlin, corporate member of Freie Universität Berlin and Humboldt-Universität zu Berlin, Charitéplatz 1, 10117 Berlin, Germany; Charité – Universitätsmedizin Berlin, corporate member of Freie Universität Berlin and Humboldt-Universität zu Berlin, NeuroCure Cluster of Excellence, 10117 Berlin, Germany; Max Delbrück Center for Molecular Medicine in the Helmholtz Association, Robert-Rössle-Straße 10, 13125 Berlin, Germany; Center for Synaptic Neuroscience and Technology, Istituto Italiano di Tecnologia, 16163 Genoa, Italy; IRCCS Ospedale Policlinico San Martino, 16132 Genoa, Italy; German Center for Neurodegenerative Diseases (DZNE) Berlin, 10117 Berlin, Germany; Charité – Universitätsmedizin Berlin, corporate member of Freie Universität Berlin and Humboldt-Universität Berlin, Einstein Center for Neurosciences, 10117 Berlin, Germany; Humboldt-Universität zu Berlin, Bernstein Center for Computational Neuroscience, Philippstraße 13, 10115 Berlin, Germany

## Abstract

Synapsins are highly abundant presynaptic proteins that play a crucial role in neurotransmission and plasticity via the clustering of synaptic vesicles. The synapsin III isoform is usually downregulated after development, but in hippocampal mossy fiber *boutons* it persists in adulthood. Mossy fiber *boutons* express presynaptic forms of short- and long-term plasticity, which are thought to underlie different forms of learning. Previous research on synapsins at this synapse focused on synapsin isoforms I and II. Thus, a complete picture regarding the role of synapsins in mossy fiber plasticity is still missing. Here, we investigated presynaptic plasticity at hippocampal mossy fiber *boutons* by combining electrophysiological field recordings and transmission electron microscopy in a mouse model lacking all synapsin isoforms. We found decreased short-term plasticity - i.e. decreased facilitation and post-tetanic potentiation - but increased long-term potentiation in male synapsin triple knockout mice. At the ultrastructural level, we observed more dispersed vesicles and a higher density of active zones in mossy fiber *boutons* from knockout animals. Our results indicate that all synapsin isoforms, including synapsin III, are required for fine regulation of short- and long-term presynaptic plasticity at the mossy fiber synapse.

**Significance statement:** Synapsins cluster vesicles at presynaptic terminals and shape presynaptic plasticity at giant hippocampal mossy fiber *boutons*. Deletion of all synapsin isoforms results in decreased short- but increased long-term plasticity.

## Introduction

Neurotransmission is a fundamental process that enables us to sense the world around us, to react to it, to think, learn and remember. This process requires high temporal and spatial fidelity, and the energy-expensive and complex regulation of synaptic vesicle trafficking is a prerequisite. A crucial aspect is the spatial arrangement of neurotransmitter-filled vesicles inside the synapse, regulated by the protein family of synapsins (Atias et al., 2019; Sansevrino et al., 2023).

Synapsins are highly abundant phosphoproteins associated with the surface of synaptic vesicles (De Camilli et al., 1990; Cesca et al., 2010), encoded by three mammalian genes (*SYN1, SYN2, SYN3*) (Südhof et al., 1989; Kao et al., 1998). Impairment of synapsin I (SynI) and II (SynII) causes vesicle dispersion and shrinks the distal vesicle cluster, the reserve pool (Li et al., 1995; Pieribone et al., 1995; Rosahl et al., 1995). Thus, synapsins main function is to control mobilization from the reserve pool, in a phosphorylation-dependent manner (Sihra et al., 1989; Hosaka et al., 1999; Chi et al., 2001). How synapsins preserve this pool is still under debate. Likely mechanisms are: (1) synapsins crosslink the vesicles, acting as tethers (Hirokawa et al., 1989), (2) synapsins form a liquid phase, capturing vesicles in it (Milovanovic et al., 2018; Pechstein et al., 2020) or (3) a mixture of both, since these mechanisms are not mutually exclusive (Zhang and Augustine, 2021; Song and Augustine, 2023).

While SynI and SynII are expressed in mature synapses (De Camilli et al., 1983; Browning et al., 1987), synapsin III (SynIII) is primarily expressed during development: after one week postnatal its levels decrease drastically (Ferreira et al., 2000) and remain low in adults (Kao et al., 1998). However, in brain regions featuring postnatal neurogenesis, SynIII is still expressed in adult tissue (Pieribone et al., 2002). This includes the dentate gyrus and hippocampal mossy fibers.

Hippocampal mossy fibers are thought to be involved in learning, memory and spatial navigation (Rolls, 2018). They connect granule cells and CA3 pyramidal cells via mossy fiber *boutons*, highly-plastic synapses (Nicoll and Schmitz, 2005). Activity-dependent changes in neurotransmission can be studied very well in these *boutons*, because they can react to a wide range of frequencies (Salin et al., 1996) and express presynaptic short- and long-term potentiation (STP, LTP) (Zalutsky and Nicoll, 1990; Nicoll and Schmitz, 2005). Recently, a mechanism for short-term memory has been proposed: the formation of a “pool engram” – an increased readily releasable pool (RRP) – which could depend on the vesicle mobilization via synapsins (Vandael et al., 2020). Unlike STP, mossy fiber LTP is still more enigmatic: It is known to be protein kinase A (PKA)-dependent (Weisskopf et al., 1994), but the precise downstream targets and potential parallel mechanisms are not yet clarified (Monday et al., 2018, 2022; Shahoha et al., 2022).

Synapsin-dependent mossy fiber physiology has been investigated in SynI/SynII double knockout (SynDKO) animals (Spillane et al., 1995; Owe et al., 2009): field recordings revealed impaired frequency facilitation in physiologically relevant ranges (Owe et al., 2009), while LTP was unchanged (Spillane et al., 1995). However, enrichment of SynIII close to the active zone at mossy fiber *boutons* (Owe et al., 2009) raised the question, if the additional knockout (KO) of SynIII would have further effects on mossy fiber transmission and plasticity.

Here, we examined a glutamatergic synapse that retains SynIII expression in adulthood and asked how neurotransmission is changed upon the complete loss of synapsins. We investigated this question in acute slices of SynI/SynII/SynIII triple knockout (SynTKO) male mice using a combined approach of transmission electron microscopy (TEM) and electrophysiological field recordings. We observed fewer vesicles in the reserve pool and increased active zone density. Field recordings provided evidence that synapsins are crucial for both STP and LTP in mossy fibers: facilitation and post-tetanic potentiation were impaired, while LTP was enhanced.

## Methods

### Reporting guidelines

This study was reported in accordance with the SAGER guidelines (Heidari et al., 2016) and ARRIVE guidelines 2.0 (Percie du Sert et al., 2020). The checklist for the SAGER guideline is provided in table 1, the checklist for the essential ten of the ARRIVE guideline is provided in table 2 and the checklist for the recommended set of the ARRIVE guideline is provided in table 3.

**Table 1:** SAGER guidelines checklist – other studies (applied sciences, cell biology, etc.). Adapted from Heidari et al., 2016. *These points extend beyond the original SAGER table.

| Section/<br>Topic | Item<br>number | Checklist item | Reported on<br>page number |
| --- | --- | --- | --- |
| General | 1 | The terms sex/gender used appropriately |  |
| Title | 2a | Title specifies the sex of animals or any cells, tissues, and other material derived from these | 1 |
|  | 2b | In applied sciences (technology, engineering, etc.), the title indicates if the study model was based on one sex/gender or the application was considered for the use of one specific sex/gender |  |
| Abstract | 3a | Abstract specifies sex of animals or any cells, tissues, and other material derived from these | 4 |
|  | 3b | In applied sciences (technology, engineering, etc.), the abstract indicates if the study model was based on one sex/gender or the application was considered for the use of one specific sex/gender |  |
| Introduction | 4a | If relevant, previous studies that show presence or lack of sex or gender differences or similarities are cited |  |
|  | 4b | Mention of whether sex/gender might be an important variant and if differences might be expected |  |
| Methods | 5a | In cell biological, molecular biological, or biochemical experiments, the origin and sex chromosome constitutions of cells or tissue cultures are stated. If unknown, the reasons are stated |  |
|  | 5b | For studies testing devices or technology, explanation of whether the product will be applied or used by all genders and if it has been tested with a user's gender in mind |  |
|  | 5c | If relevant, description of how sex/gender was considered in the design |  |
|  | 5d | For in-vivo and in-vitro studies using primary cultures of cells, or cell lines from humans or animals, or ex-vivo studies with tissues from humans or animals, the sex of the subjects or source donors is stated (except for immortalized cell lines, which are highly transformed) | 7 |
| Results | 6 | For studies using animal models, present a sex breakdown of the animals* |  |
| Discussion | 7 | If relevant, potential implications of sex/gender on the study results and analyses, including the extent to which the findings can be generalized to all sexes/genders in a population | 26 |

**Table 2:**
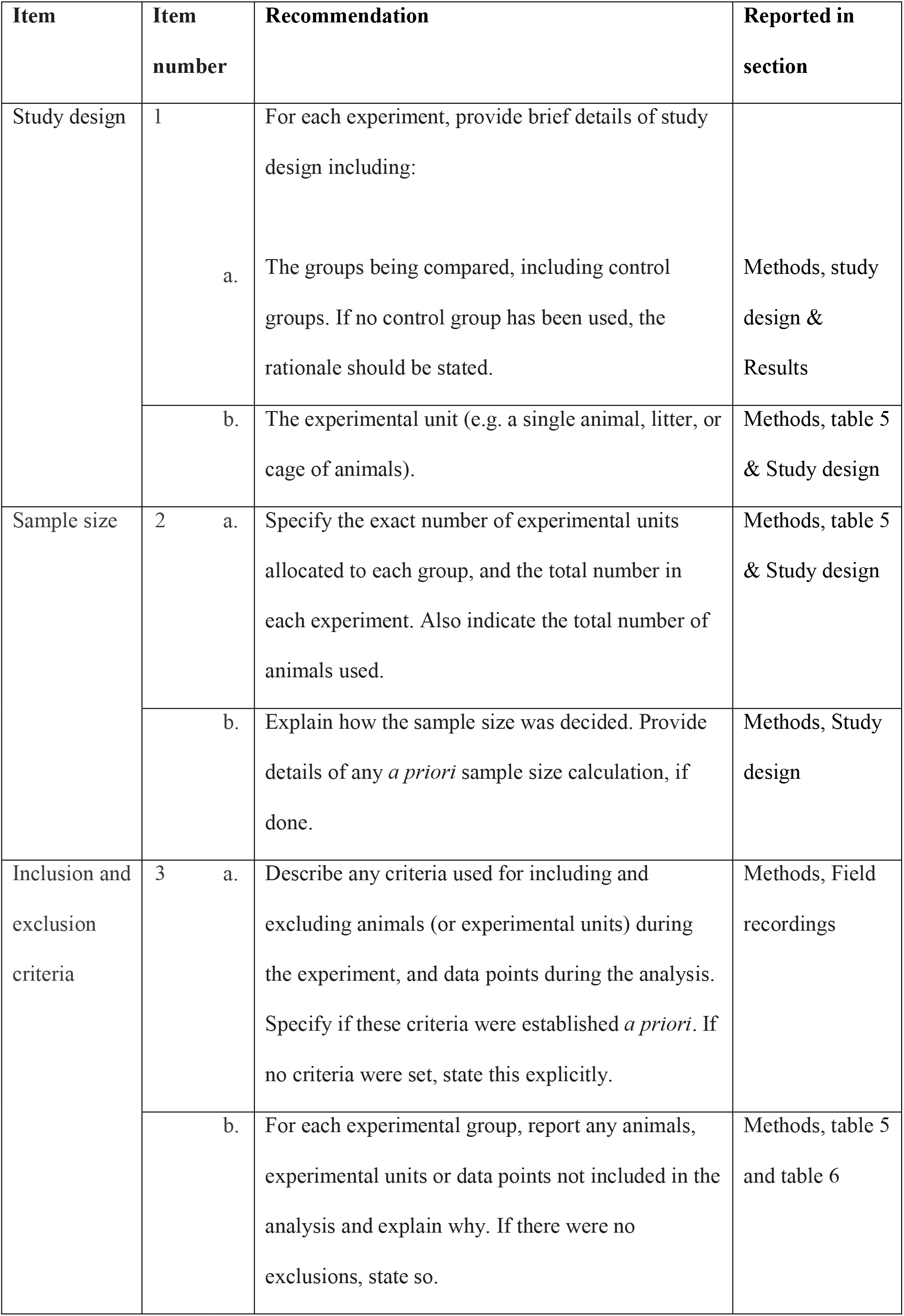
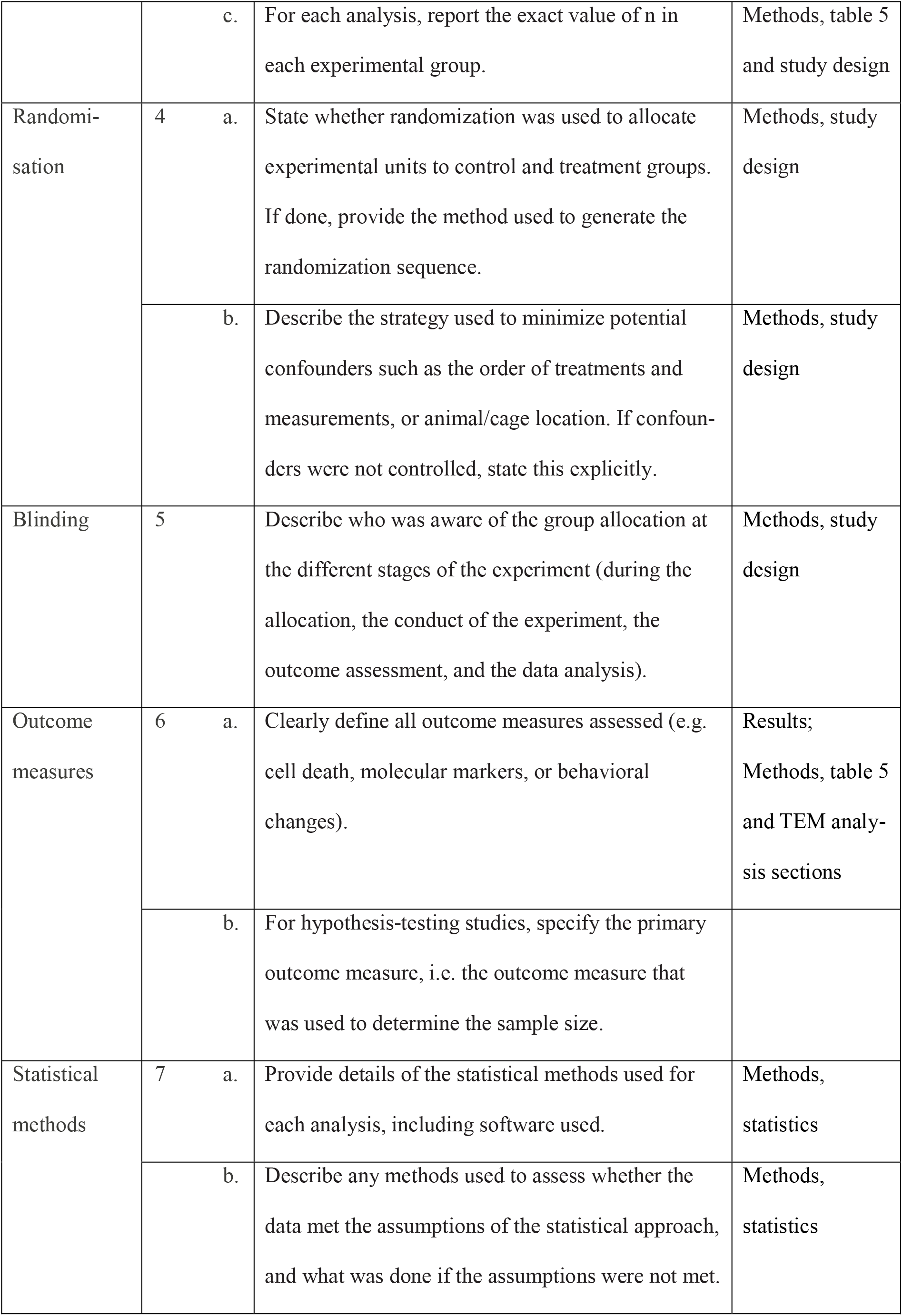
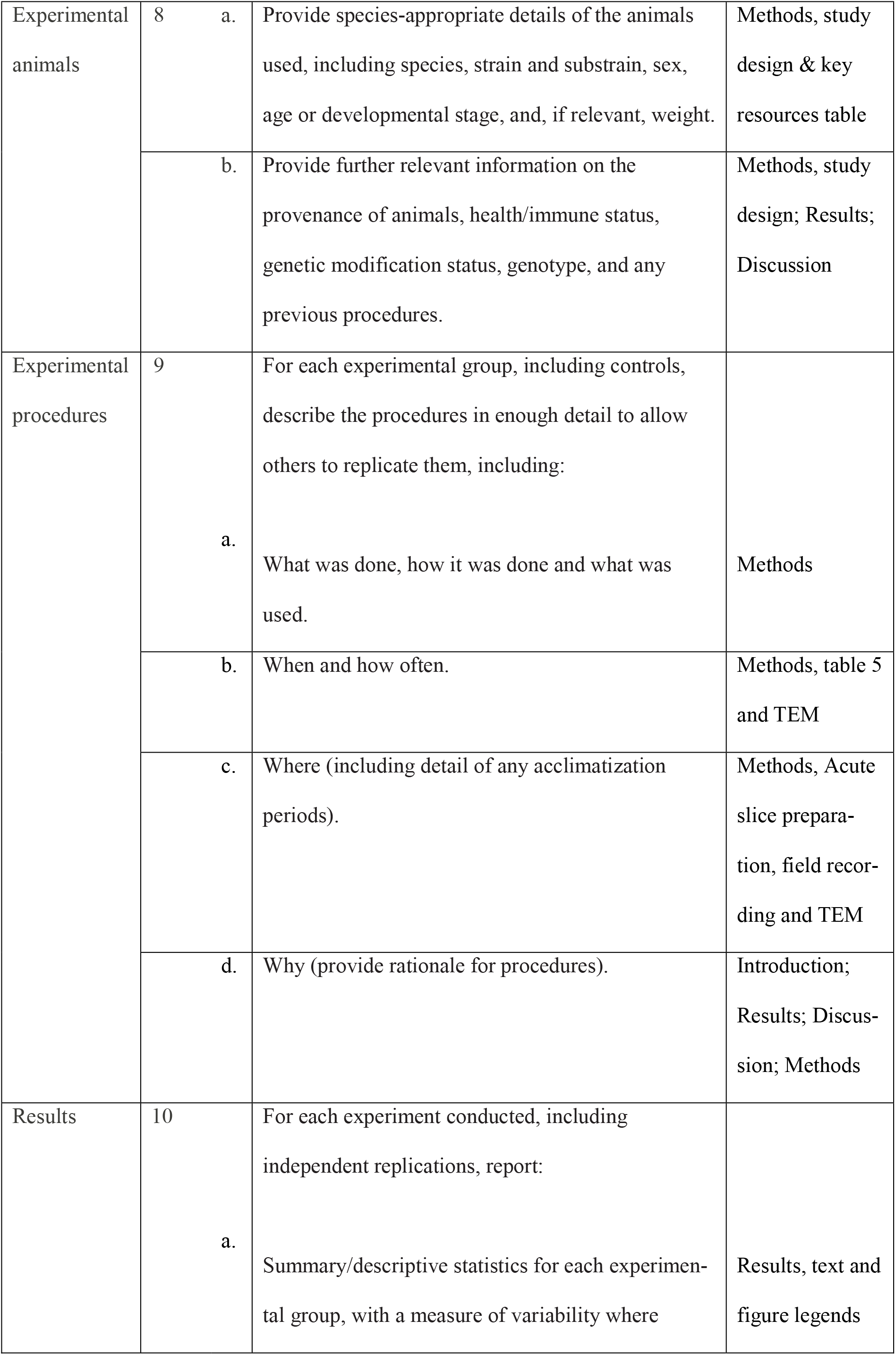
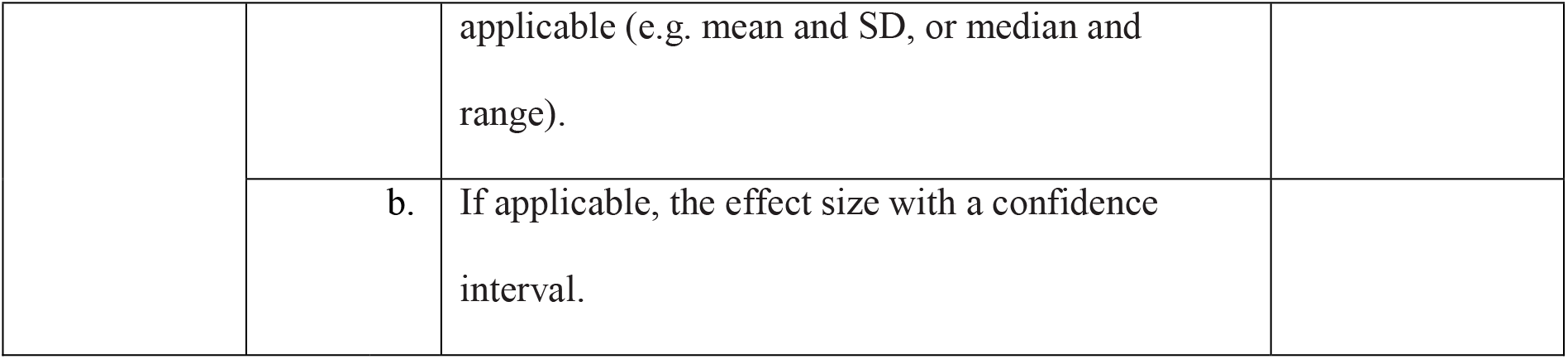
The ARRIVE guidelines 2.0 checklist: the essential ten. Adapted from Percie du Sert et al., 2020.

**Table 3:**
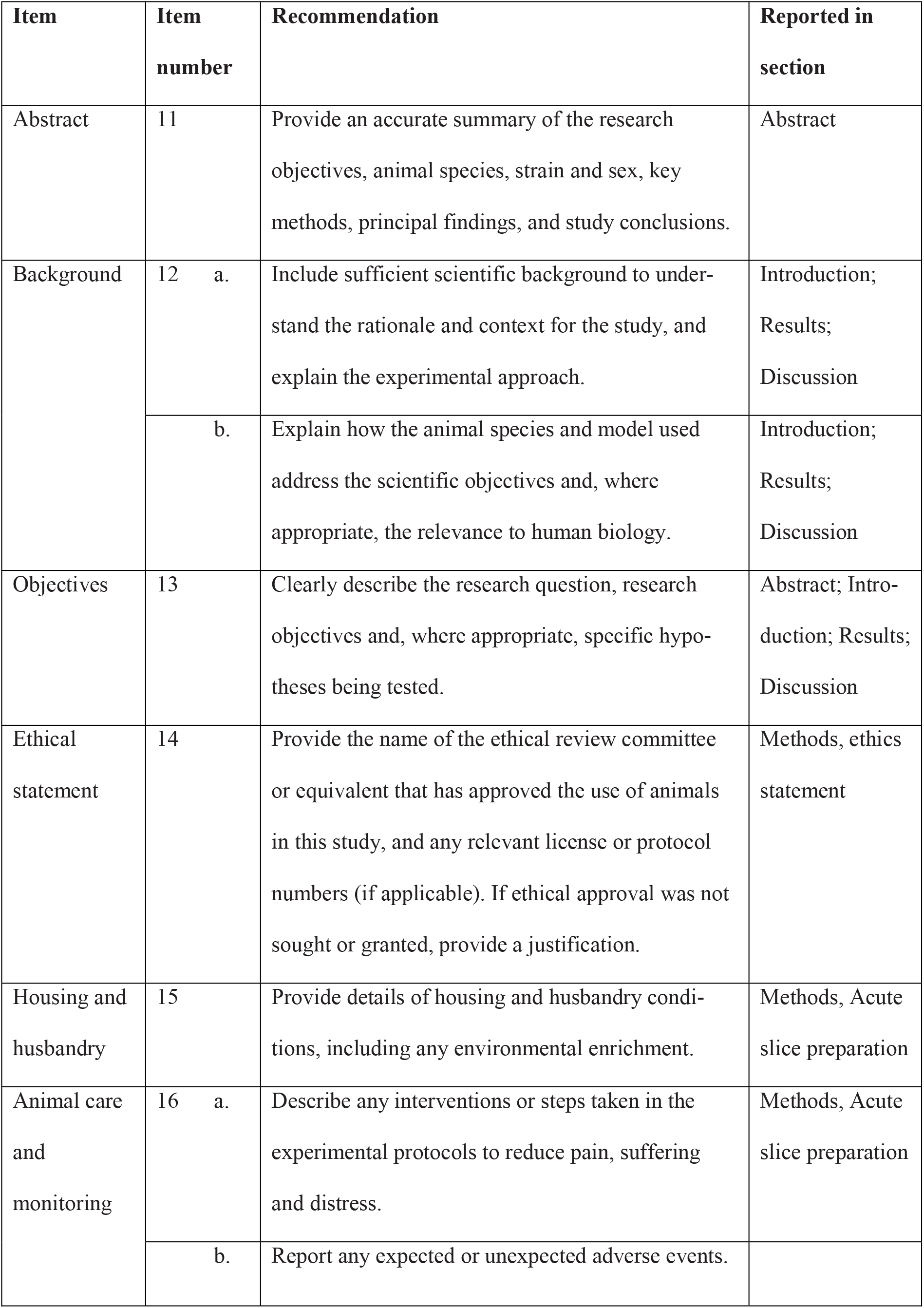
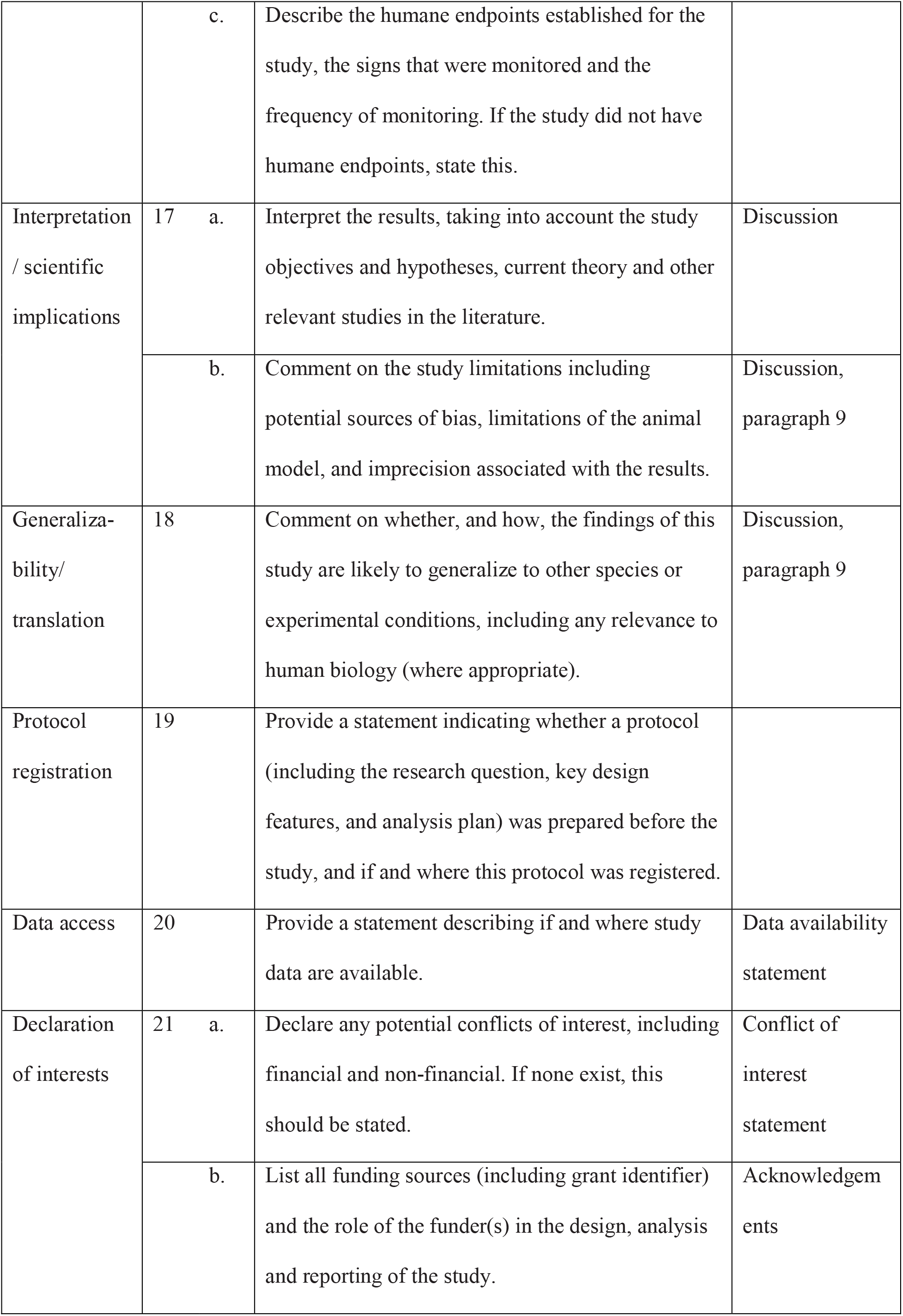
The ARRIVE guidelines 2.0: the recommended set. Adapted from Percie du Sert et al., 2020.

| Item | Item number | Recommendation | Reported in section |
| --- | --- | --- | --- |
| Abstract | 11 | Provide an accurate summary of the research objectives, animal species, strain and sex, key methods, principal findings, and study conclusions. | Abstract |
| Background | 12 a. | Include sufficient scientific background to understand the rationale and context for the study, and explain the experimental approach. | Introduction;<br>Results;<br>Discussion |
|  | b. | Explain how the animal species and model used address the scientific objectives and, where appropriate, the relevance to human biology. | Introduction;<br>Results;<br>Discussion |
| Objectives | 13 | Clearly describe the research question, research objectives and, where appropriate, specific hypotheses being tested. | Abstract; Introduction; Results;<br>Discussion |
| Ethical statement | 14 | Provide the name of the ethical review committee or equivalent that has approved the use of animals in this study, and any relevant license or protocol numbers (if applicable). If ethical approval was not sought or granted, provide a justification. | Methods, ethics statement |
| Housing and husbandry | 15 | Provide details of housing and husbandry conditions, including any environmental enrichment. | Methods, Acute slice preparation |
| Animal care and monitoring | 16 a. | Describe any interventions or steps taken in the experimental protocols to reduce pain, suffering and distress. | Methods, Acute slice preparation |
|  | b. | Report any expected or unexpected adverse events. |  |

|  |  |  |  |  |
| --- | --- | --- | --- | --- |
|  |  | c. | Describe the humane endpoints established for the study, the signs that were monitored and the frequency of monitoring. If the study did not have humane endpoints, state this. |  |
| Interpretation / scientific implications | 17 | a. | Interpret the results, taking into account the study objectives and hypotheses, current theory and other relevant studies in the literature. | Discussion |
|  |  | b. | Comment on the study limitations including potential sources of bias, limitations of the animal model, and imprecision associated with the results. | Discussion, paragraph 9 |
| Generalizability/ translation | 18 |  | Comment on whether, and how, the findings of this study are likely to generalize to other species or experimental conditions, including any relevance to human biology (where appropriate). | Discussion, paragraph 9 |
| Protocol registration | 19 |  | Provide a statement indicating whether a protocol (including the research question, key design features, and analysis plan) was prepared before the study, and if and where this protocol was registered. |  |
| Data access | 20 |  | Provide a statement describing if and where study data are available. | Data availability statement |
| Declaration of interests | 21 | a. | Declare any potential conflicts of interest, including financial and non-financial. If none exist, this should be stated. | Conflict of interest statement |
|  |  | b. | List all funding sources (including grant identifier) and the role of the funder(s) in the design, analysis and reporting of the study. | Acknowledgements |

### Ethics statement

All animal experiments were carried out according to the guidelines stated in Directive 2010/63/EU of the European Parliament on the protection of animals used for scientific purposes and were approved by the animal welfare committee of Charité – Universitätsmedizin Berlin and the Landesamt für Gesundheit und Soziales (LaGeSo) Berlin (permit T 0100/03 and permit G 0146/20).

### Study design

In this study, only male mice were used for experiments to exclude possible indirect estrogen effects on mossy fiber plasticity (Harte-Hargrove et al., 2013). In electrophysiological recordings, C57BL/6J control mice (RRID:IMSR_JAX:000664) were compared to SynI/SynII/SynIII triple knockout (SynTKO) mice (RRID:MMRRC_041434-JAX) in two age groups: one younger group (4-6 weeks of age), which is referred to as presymptomatic, and one older group (17-19 weeks of age), which is referred to as symptomatic. These terms describe the phenotype before and after the onset of epileptic seizures in SynTKO animals, respectively (Farisello et al., 2013). SynTKO mice were purchased from the Jackson Laboratory (RRID:SCR_004633) and were based on work from Gitler and coworkers (Gitler et al., 2004). The presymptomatic SynTKO data were obtained from two different cohorts. We received the first cohort from Prof. Dr. Fabio Benfenati (Instituto Italiano di Tecnologia, Genova, Italy). The second cohort from Dr. Dragomir Milovanovic (DZNE, Berlin, Germany) was housed and bred in the Charité animal facility (FEM; Forschungseinrichtungen für Experimentelle Medizin). Symptomatic SynTKO animals and all control animals were also bred and born in the Charité animal facility. For each experiment we were aiming for at least three biological replicates (animals) per group. Depending on experimental success (how many recordings needed to be excluded, technical failures), we added more animals per group.

### Field recordings

Data from both presymptomatic SynTKO cohorts were pooled, because they were not significantly different (Table 4). Field recording experiments in all four groups (WT, SynTKO, presymptomatic, symptomatic) were repeated with at least three mice from more than one litter (Table 5). Variable *s* represents the number of recorded slices while *a* reports the number of animals. We were not blinded towards the genotype, because the phenotype was too strong.

**Table 4:**
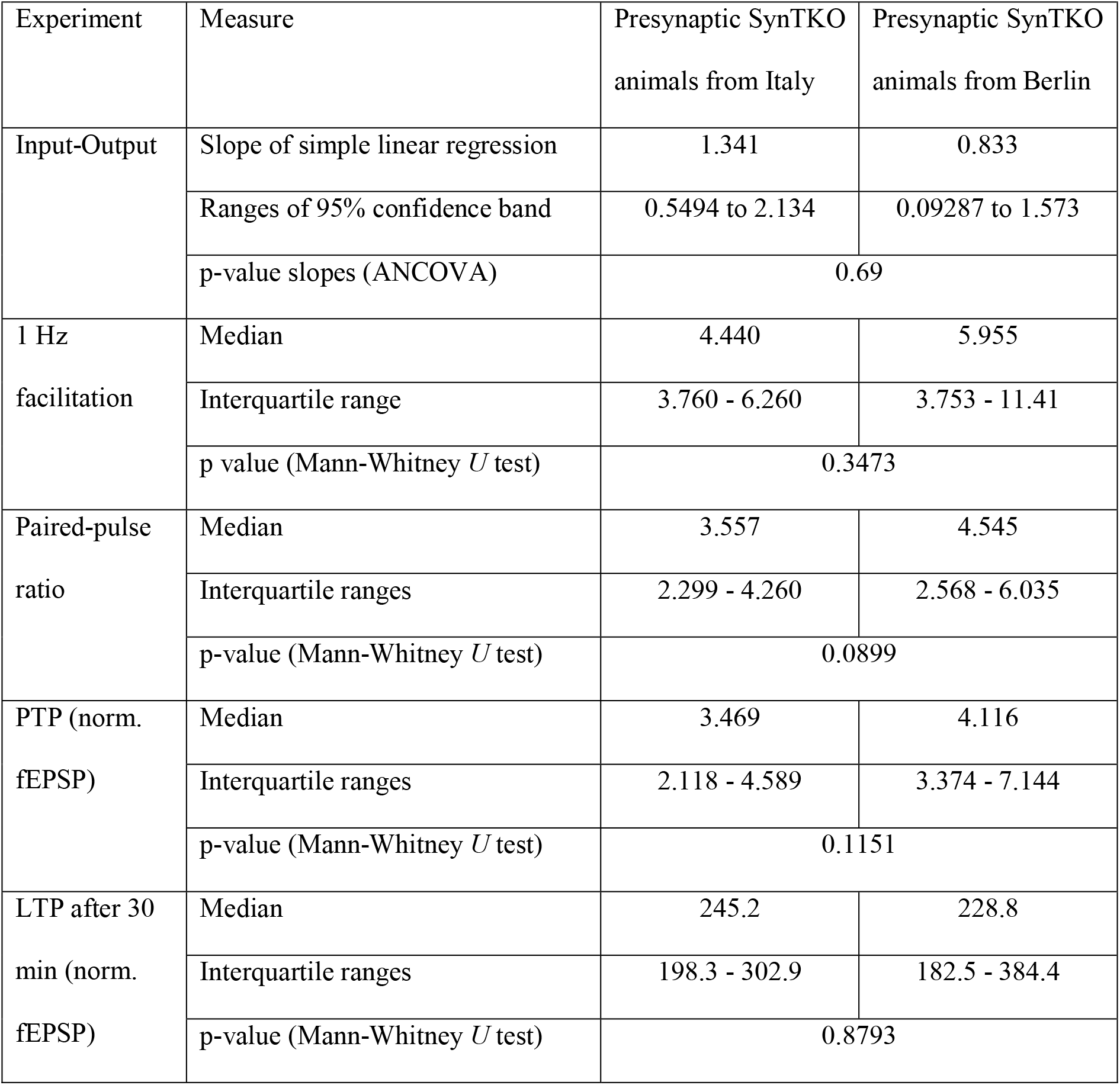
Statistical comparison for experimental values between two cohorts of presymptomatic SynTKO animals.

**Table 5:**
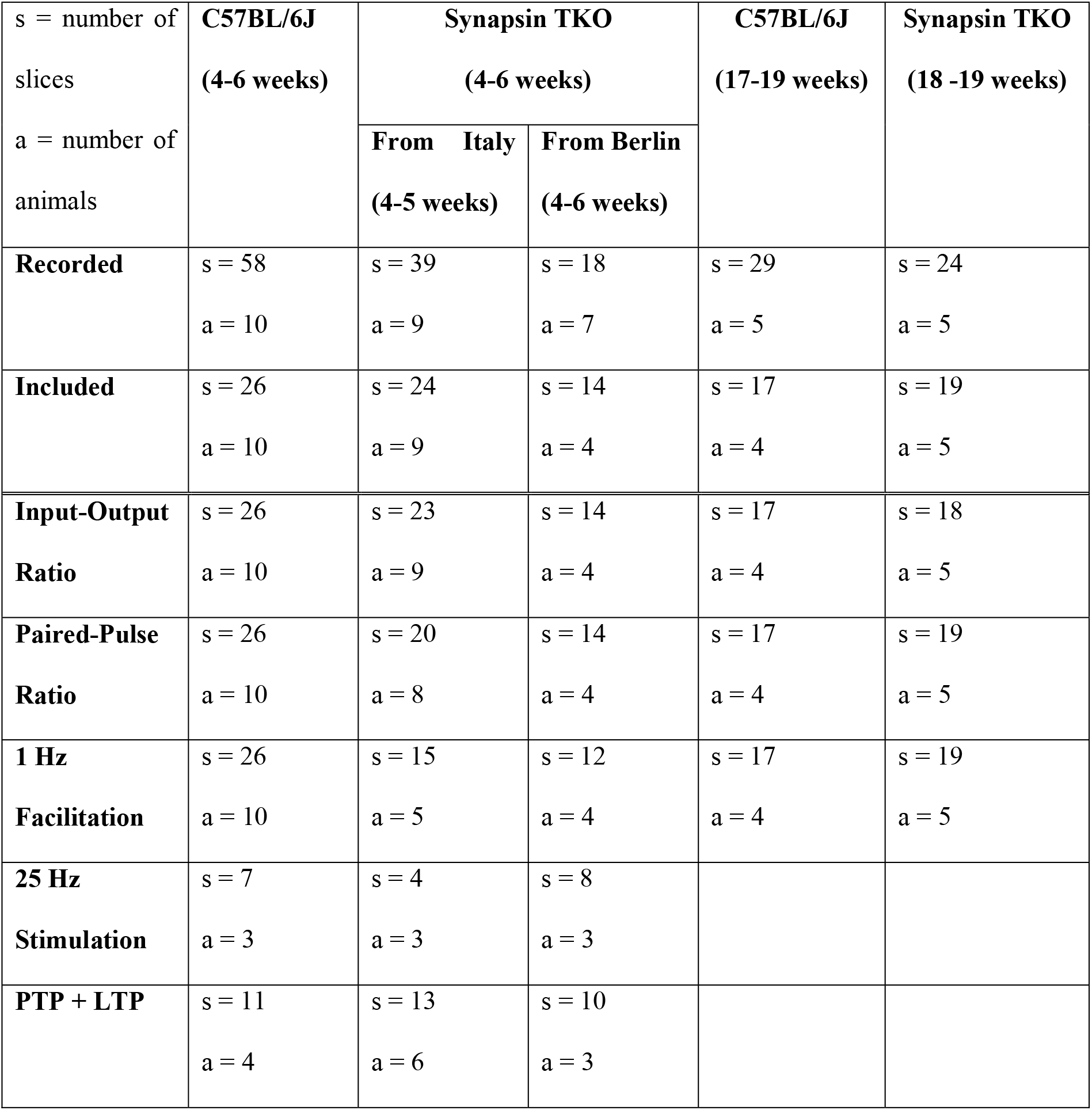
Overview of slice and animal numbers for different experimental groups for field recordings. Note: all numbers reported for individual experiments are only from the included subset of recordings.

Recordings were excluded when they had a baseline fEPSP smaller than two times noise (Table 6). Noise was approximately 25 µV, so the baseline fEPSP amplitude needed to be at least 50 µV to be included. Furthermore, to include only mossy fiber specific recordings, we applied 1 µM DCG-IV (#0975, Tocris Bioscience) at the end of each experiment (Kamiya et al., 1996). If the suppression was 75% or more, the recording was included (Table 6). We were not able to measure input-output curves for all animals. For those cases where it was not recorded with different input strengths, we took the averaged baseline values for PFV and fEPSP, respectively. If the PFV could not be measured unambiguously, this measurement was excluded from the input-output graph. If the 1 Hz or 25 Hz induction failed, the respective measurements were excluded for analysis, but all other parameters from the same experiment were included. The same was true for some recordings, in which no 25 Hz stimulation and thus no PTP and LTP recordings were conducted. If possible, two mice with different genetic backgrounds were recorded on the same day to minimize variability due to experimental day.

**Table 6:** Exclusion reasons for field recordings. Note that several reasons can apply to the same recording.

| **Excluded recordings** | **C57BL/6J**<br>**(4-6 weeks)** | **Synapsin**<br>**TKO**<br>**(4-6 weeks)** | **C57BL/6J**<br>**(17-19 weeks)** | **Synapsin TKO**<br>**(18 -19 weeks)** |
| --- | --- | --- | --- | --- |
| **Total number** | 32 | 19 | 12 | 5 |
| **Baseline EPSP < 50 μV** | 4 | 2 | 2 | 2 |
| **DCG-IV effect < 75%** | 25 | 14 | 11 | 4 |
| **Other reasons** | 4 | 2 | 0 | 1 |

### Transmission electron microscopy

For ultrastructural investigation of mossy fiber *boutons* male mice at the age of 4-6 weeks were used. Data from the two presymptomatic SynTKO cohorts were pooled. For vesicle numbers and mean nearest neighbor distance we identified mossy fiber *boutons* from two young WT and three SynTKO mice. We imaged serial sections from 12 WT and 16 SynTKO mossy fiber *boutons*, respectively. For active zone density we analyzed partial 3D reconstructions of mossy fiber *boutons* from three WT and three SynTKO animals. Slices from each animal were either treated with forskolin or allocated as control. Allocation of slices to treatment or control group was block-randomized. Replicates of 17 (WT), 16 (WT + forskolin), 16 (SynTKO) and 18 (SynTKO + forskolin) mossy fiber *boutons* were analyzed. Number n represents the number of presynaptic *bouton* reconstructions. The experimenter was blinded to the treatment of slices from fixation of the slices until the end of analysis. Due to the strong reduction in vesicle density of SynTKO synapses, blinding during analysis was only possible between treatment groups, but not between genotypes.

### Acute slice preparation

Animals were kept in a 12L:12D hour light-dark cycle and water and food were provided *ad libitum*. Cages offered shelter in form of a house and tubes. Cages of SynTKO animals were kept in remote shelves to minimize exposure to light and possible noises. The first cohort of presymptomatic SynTKO animals was imported from Italy and allowed to sit in the Charité animal facility for several days before the experiments started. After transfer from the animal facility to the preparation room, all animals were allowed to acclimate to the new surrounding for at least half an hour. Acute brain slices were prepared as follows: Mice were anaesthetized under the hood with isoflurane and quickly sacrificed with sharp scissors. The brain was taken out and placed in oxygenated ice-cold sucrose-artificial cerebrospinal fluid (S-ACSF) for three minutes to allow equilibration. S-ACSF contained in mM: 50 NaCl, 25 NaHCO_3_, 10 Glucose, 150 Sucrose, 2.5 KCl, 1 NaH_2_PO_4_, 0.5 CaCl_2_, 7 MgCl_2_. All solutions were saturated with 95% O_2_ (vol/vol) / 5% CO_2_ (vol/vol) and had a pH of 7.4 and an osmolarity of 340 mOsm. Hemispheres were separated and 300 µm/150 µm (field recordings/electron microscopy) thick sagittal sections were cut from both hemispheres with a vibratome (VT1200 S, Leica Biosystems (RRID:SCR_018453)). Slices were stored in a submerged chamber in oxygenated S-ACSF at 34°C for half an hour before they were moved to another submerged chamber with artificial cerebrospinal fluid (ACSF) at room temperature. There, slices were kept until the start of experiments. ACSF had an osmolarity of 300 mOsm and a pH of 7.4 and contained the following substances in mM: 119 NaCl, 26 NaHCO_3_, 10 Glucose, 2.5 KCl, 1 NaH_2_PO_4_, 2.5 CaCl_2_, 1.3 MgCl_2_. All chemicals were purchased from Sigma Aldrich.

### Field recordings

Slices were kept in a submerged chamber with ACSF at least 30 minutes and up to six hours before start of recordings. Slices were placed in a recording chamber under a microscope and were continuously superfused with oxygenated ACSF at room temperature at a rate of approximately 2.5 ml/min. The recording electrode was fixed in a headstage of the amplification system (Axon Instruments, MultiClamp 700A/700B (RRID:SCR_018455)). Stimulation and recording electrode units were placed on micromanipulators (Mini 23/25, Luigs & Neumann GmbH) for precise movement control via a control system (SM-5/-7/-10, Luigs & Neumann GmbH).

The stimulation and recording electrodes were prepared from silver wires (AG-8W and E-205, Science Products). Glass pipettes were made from borosilicate capillaries (GB150EFT-10, Science Products or #1403005, Hilgenberg) with a pipette puller (PC-10, Narishige or DMZ-Universal Puller, Zeitz-Instrumente) and were broken at the tip with a micro forge (MF-830, Narishige) to receive low-resistance pipettes. Electrodes were placed in the hilus of the dentate gyrus near the granule cell layer (stimulation) and within the *stratum lucidum* of the area CA3 of the hippocampus (recording), respectively.

Stimulations were executed with a stimulation box (ISO-Flex, A.M.P.I. (RRID:SCR_018945)) and stimulation patterns were controlled with a Master 8 generator (A.M.P.I. (RRID:SCR_018889)). Igor Pro (version 6, WaveMetrics (RRID:SCR_000325)) was used for signal acquisition. The Axon MultiClamp amplifier (700A/700B, Molecular Devices (RRID:SCR_018455)) was used in current clamp mode I=0, with filtering of 2 kHz. Signals were digitized (Axon Digidata 1550B, Molecular Devices / BNC-2090; National Instruments Germany GmbH) at a rate of 20 kHz. Mossy fiber signals were searched by placing the stimulation and recording electrodes at different locations in the hilus and *stratum lucidum*, respectively. Once a mossy fiber input was obtained, the recording was started and the mossy fibers were stimulated at 0.05 Hz.

The standard stimulation frequency was 0.05 Hz throughout the experiment, unless otherwise stated. Recorded sweep length was 0.5 s except for the high-frequency stimulation at 25 Hz where 5.5 s were recorded. First, input-output relations were recorded by applying different input currents via the stimulation box. The strength of the input current was adjusted to yield a specific presynaptic fiber volley size: 0.05 mV, 0.1 mV, 0.2 mV, 0.3 mV and maximum (maximal stimulation strength of 10 mA). Each input strength was recorded for three sweeps. Afterwards, a medium stimulation strength was chosen and a baseline was recorded for at least ten sweeps. Then, the stimulation frequency was increased to 1 Hz for 20 sweeps for recording of frequency facilitation. Afterwards, when fEPSP amplitudes declined to baseline level again, a paired-pulse with an inter-stimulus interval of 50 ms was applied for three sweeps. Then, a baseline was recorded for 10 minutes (30 sweeps, except for once when only 20 sweeps were recorded) before a high-frequency train of stimuli was given: four times 125 pulses at 25 Hz every 20 seconds with a recorded sweep length of 5.5 seconds. Post-tetanic potentiation and subsequently long-term potentiation were measured for at least 30 minutes after the tetanus. Mossy fiber purity of signals was verified at the end of each recording with the application of 1 µM DCG-IV (#0975, Tocris Bioscience). All recordings with a suppression of at least 75% of the signal were used for analysis.

### Field recording analysis

Field recordings were analyzed with Igor Pro (versions 6 and 8, WaveMetrics (RRID:SCR_000325)) and the installed plugin NeuroMatic (RRID:SCR_004186) as well as Microsoft Excel (RRID:SCR_016137). Igor Pro is commercially available at https://www.wavemetrics.com/products/igorpro and Microsoft Excel is commercially available at https://www.microsoft.com/de-de/microsoft-365/excel. Presynaptic fiber volleys (PFV) were measured peak to peak. Field EPSP amplitudes were baseline-corrected and measured +/- 2 ms around the peak. For input-output curves, the mean value of the three sweeps at the same stimulation strength was taken, except for the cases in which no input-output curve was recorded: here, we took the average size of PFV and fEPSP amplitude from the initial baseline. Field EPSP amplitudes during 1 Hz facilitation were normalized to the initial baseline (10 sweeps, 3 minutes). The paired-pulse ratio (PPR) was calculated as the ratio between the second to the first fEPSP amplitude. The stated PPR refers to the first of three paired stimulations. For analysis of the high-frequency trains we normalized the fEPSP amplitudes to the baseline before (30 sweeps, 10 minutes). We also evaluated the PFV size for a subset of fEPSPs of the 25 Hz trains. We measured the PFV for stimuli 10-15 and averaged those six values for the first and fourth stimulation train, respectively (Figure 2-1b). Also, we calculated the ratio of those averaged values between fourth and first stimulation train, to compare the relative loss of PFV size (Figure 2-1c). Values for PTP and LTP were normalized to the average of the recorded baseline before high-frequency stimulation (30 sweeps, 10 minutes). Values for LTP were the averaged fEPSP amplitudes from minute 20-30 (30 sweeps) after induction. At the end of the recording, specificity was verified by application of DCG-IV. We averaged the last 15 sweeps of DCG-IV wash-in for quantification. Recordings, in which the suppression was less than 75% were not counted as mossy fiber-specific and were not included in the analysis.

### Conventional electron microscopy

After preparation, acute slices were allowed to recover in ACSF at room temperature for at least 30 minutes. Subsequently we induced chemical LTP in half of the slices by incubating them in 50 µM forskolin (AG-CN2-0089-M050, Cayman Chemical), solved in DMSO, for 15 minutes at room temperature in oxygenated ACSF (Orlando et al., 2021). The other half of the slices (controls) were incubated in ACSF containing the same concentration of DMSO as the treatment group. Treatment was allocated following a block randomization design. Subsequently, we moved the slices under a chemical hood where fixation, post-fixation, staining, dehydratation, and infiltration steps were performed. We fixed proteins by immersing brain slices in a solution containing 1.25% glutaraldehyde (#E16216, Science Services) in 66 mM NaCacodylate (#E12300, Science Services) buffer for 1 hour at room temperature. After washes in 0.1 M NaCacodylate buffer slices were postfixed in 1% OsO_4_ in 0.1 M NaCacodylate buffer for 1 hour at room temperature. Slices were then washed and stained en bloc with 1% uranyl acetate (#1.08473, Merck) in dH_2_O and dehydrated in solutions with increasing ethanol concentration (70%, 80%, 96%, 100%). Final dehydration was obtained by incubating slices in propylene oxide (#20401, Electron Microscopy Sciences). The infiltration of epoxy resin was obtained by serial incubations in increasing resin/propylene oxide dilutions (1:3; 1:1; 3:1). Samples were finally flat embedded in Epon (#E14120-DMP, Science Services) for 48 hours at 60°C. The *stratum lucidum* in the CA3 region of the hippocampus was identified in 700 nm semi-thin sections stained with Toluidine blue (Sigma) using a light microscope (Olympus); 70 nm serial sections of these regions of interest were cut with an Ultracut UCT ultramicrotome (Leica Microsystems) equipped with an Ultra 45° diamond knife (Diatome) and collected on pioloform-coated copper slot grids (#EMS2010-Cu, Science Services). If not otherwise stated, all chemicals were purchased from EMS - Electron Microscopy Sciences and sold by Science Services.

### Electron microscopy imaging of serial sections and 3D reconstructions

Synapses were identified and imaged at 20 kx using a EM 900 Transmission Electron Microscope (Carl Zeiss, RRID:SCR_021364) operated at 80 keV and equipped with a Proscan 2K Slow-Scan CCD-Camera (Carl Zeiss). The *stratum lucidum* of the hippocampal region CA3 was easily distinguishable for the presence of big mossy fiber *boutons* and for its localization just above the pyramidal cell layer. Serial images of individual mossy fiber *boutons* were manually acquired in manually collected serial sections using the ImageSP software (TRS & SysProg) and aligned using the Midas script of the IMOD Software (RRID:SCR_003297). ImageSP software is commercially available at https://sys-prog.com/en/software-for-science/imagesp/ and IMOD software is freely available at https://bio3d.colorado.edu/imod/. Synaptic profiles were manually segmented in each image of series belonging to the same mossy fiber *bouton*. Active zones were traced in IMOD as open lines in serial projections and rendered as a meshed surface. The volume of the 3D reconstruction was calculated by creating a meshed 3D volume in IMOD.

### Synaptic vesicle analysis

To analyze synaptic vesicles, we used a machine-learning-based algorithm that we had previously developed (Imbrosci et al., 2022). It is freely available at https://github.com/Imbrosci/synaptic-vesicles-detection-extra. Briefly, we manually traced the contour of the mossy fiber *bouton* using Fiji (RRID:SCR_002285), freely available at https://fiji.sc/. This approach allowed us to obtain a measure of the *bouton* area and to create a mask over parts of the image which were not relevant for analysis. Synaptic vesicle analysis was performed automatically in a batch. From all images the number of vesicles as well as the mean nearest neighbor distance were obtained. A tutorial with a demonstration of the tool can be found online at https://www.youtube.com/watch?v=cvqIcFldVPw.

### Statistics

For statistical analysis we used GraphPad Prism software (GraphPad Prism version 8.4.0 for Windows, San Diego, California USA (RRID:SCR_002798)) and R Project for Statistical Computing (version 4.2.2, RRID:SCR_001905) in RStudio (version 2022.12.0, RRID:SCR_000432). GraphPad prism is commercially available at https://www.graphpad.com/ and R Project for Statistical Computing is freely available at https://www.r-project.org/. Data were visually inspected and tested for normality (D’Agostino and Pearson test) before evaluating them statistically, to understand if the distribution was Gaussian or non-Gaussian. Individual data points are shown as median +/- quartiles, mean values with borders of 95% confidence intervals or as mean +/- SEM.

### GraphPad Prism

For Figure 1b and Figure 1-1b, data points were fitted with a simple linear regression. Slopes of those regressions were tested with a two-tailed ANCOVA and are shown with 95% confidence bands. For Figure 1c, Figure 1-1c, Figure 2b+e and Figure 3a+c, data were tested with a mixed-effects model and a post-hoc Sidak’s test for multiple comparisons. Factors time, genotype and the interaction of both were tested. For Figure 1d, Figure 1-1d, Figure 1-2a-c, Figure 2c+f, Figure 2-1c and Figure 3d, ranks were compared with Mann-Whitney *U* tests. For data in Figure 3b we used a Kruskal-Wallis test with a post-hoc Dunn’s correction for multiple comparisons. For Figure 2-1b we used a Wilcoxon test to compare ranks.

**Figure 1:**
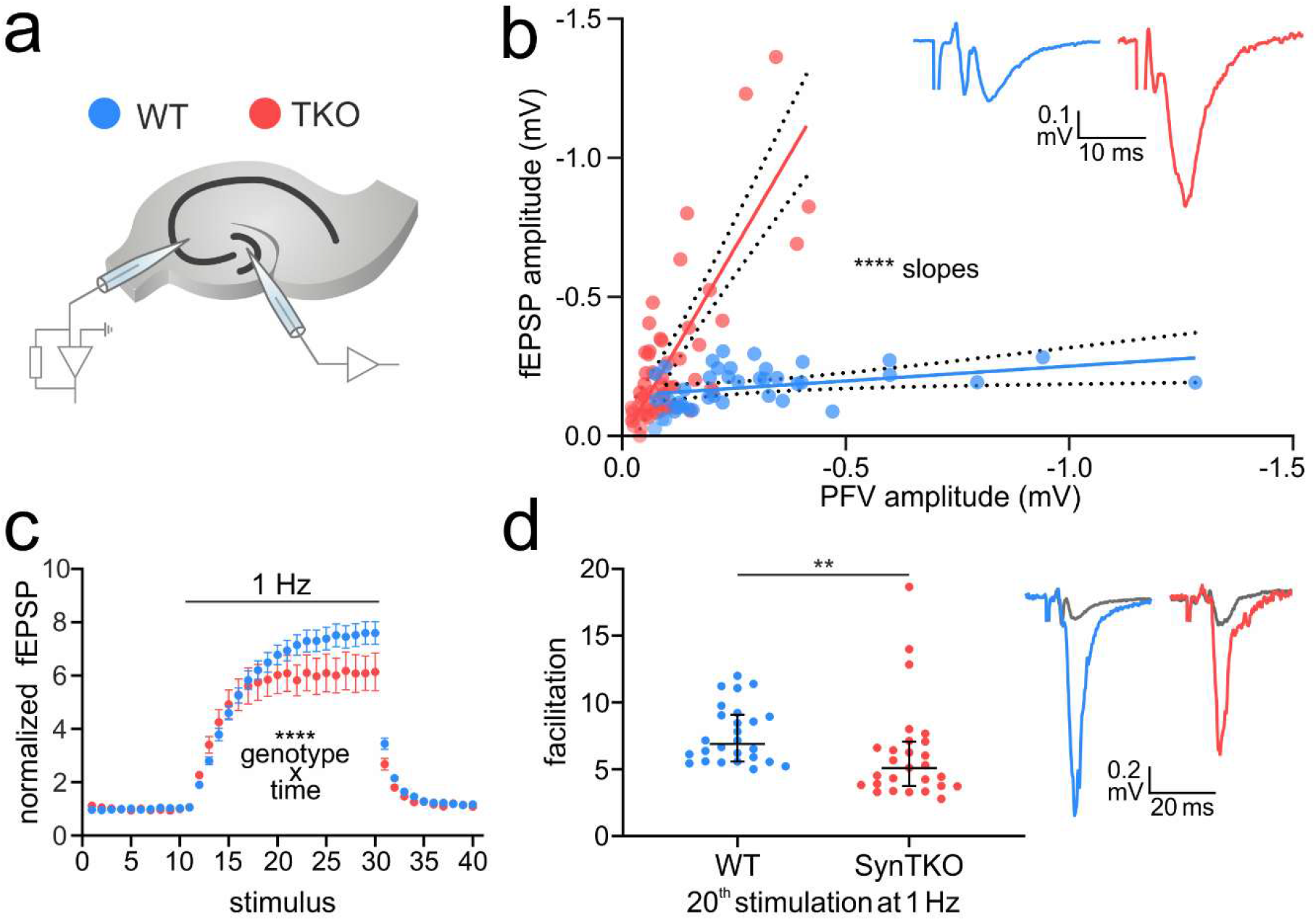
Increased excitability, but reduced facilitation, at mossy fibers of presymptomatic SynTKO mice. **a)** Field recording setup for an acute brain slice. The stimulation electrode was placed in the hilus close to the granule cell layer of the dentate gyrus, the recording electrode was placed in the stratum lucidum of area CA3 of the hippocampus where mossy fibers terminate. **b)** Excitability was increased in brain slices from SynTKO mice (red; 37 slices from 13 animals) compared to WT mice (blue; 26 slices from 10 animals). Pooled fEPSP amplitudes (mV) were plotted against pooled presynaptic fiber volley (PFV) amplitudes (mV) and fitted with a simple linear regression. The slopes of the linear regressions were significantly different (p < 0.0001, tested with a two-tailed ANCOVA). 95% confident bands are shown as dotted lines around the fit. **Inset:** Example traces from WT (blue) and SynTKO (red) slices with similar PFV amplitudes. Note the difference in the corresponding fEPSP amplitude. **c)** Frequency facilitation is reduced in SynTKO (red) compared to WT (blue) slices. Averaged normalized fEPSP amplitudes +/- SEM from all WT (blue; 26 slices from 10 animals) and SynTKO (red; 27 slices from 9 animals) recordings plotted against the number of stimuli. Stimuli 1-10 were given with a frequency of 0.05 Hz, stimuli 11-30 with 1 Hz and stimuli 31-41 with a frequency of 0.05 Hz again. Both time and the interaction between genotype and time were significantly different in a mixed-effects model (p < 0.0001). Post-hoc Sidak’s test for multiple comparisons revealed no significant differences. **d)** Facilitation was reduced in SynTKO compared to WT animals after 1 Hz frequency stimulation. **Left:** fEPSP amplitudes at the 20^th^ stimulus at 1 Hz for individual WT (blue dots; 26 slices from 10 animals) and SynTKO (red dots; 27 slices from 9 animals) recordings. Median values +/- interquartile ranges are shown in black. Facilitation was significantly different (p = 0.0029, Mann-Whitney U test). **Right:** Example fEPSP amplitudes from WT (blue) and SynTKO (red) recordings at the 20^th^ 1 Hz stimulus. Respective baseline fEPSP amplitudes are shown in grey.

**Figure 2:**
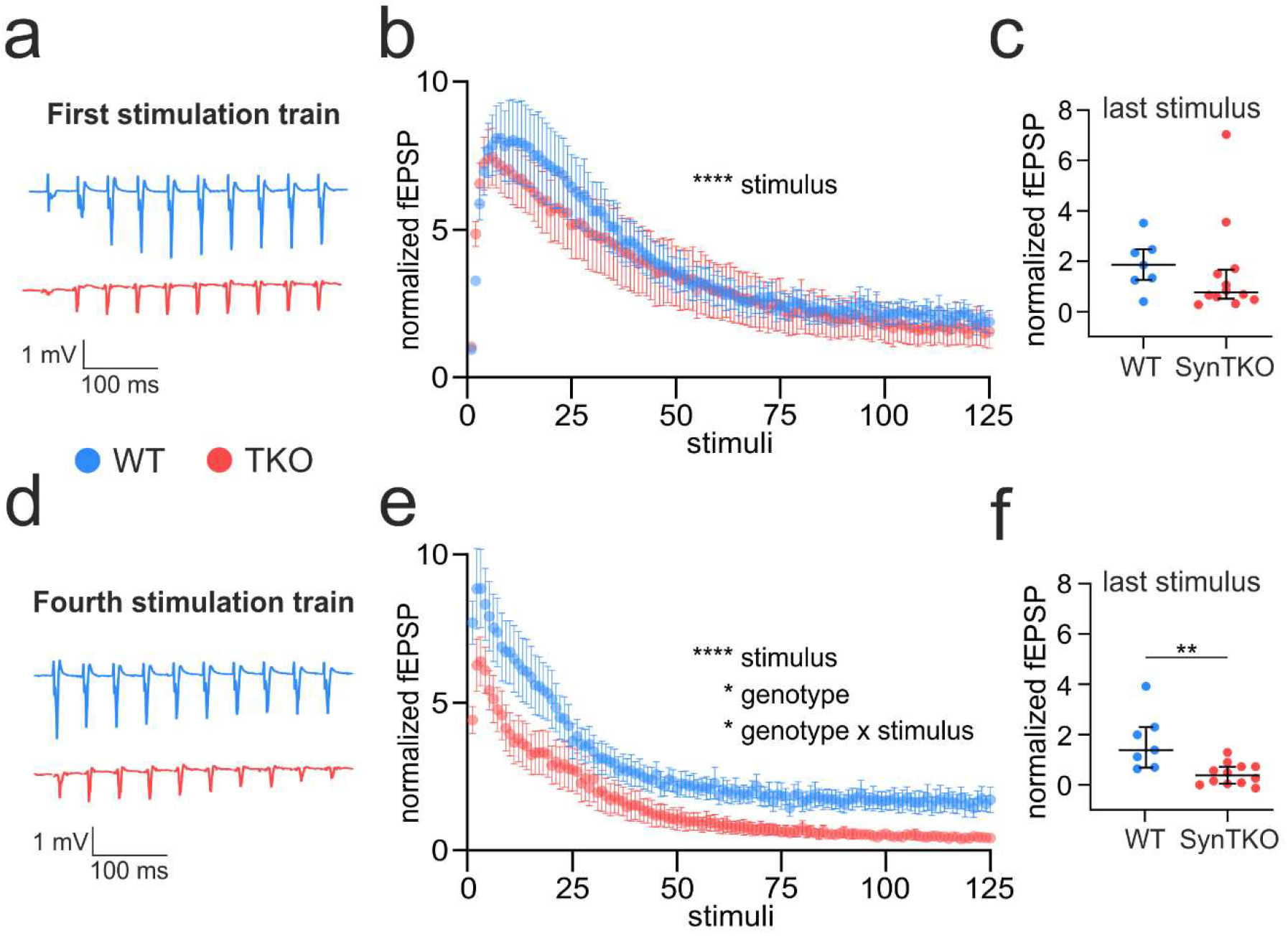
Faster depression during high-frequency stimulation in SynTKO mice. High-frequency stimulation comprised four trains of 125 pulses at 25 Hz with an interval of 20 seconds between the first stimuli of consecutive trains. **a)** Example traces show fEPSP amplitudes of mossy fibers from WT (blue) and SynTKO (red) slices in response to the first 10 stimuli of the first high-frequency stimulation train. **b)** Normalized averaged fEPSP amplitudes plotted against number of stimuli of the first high-frequency stimulation train for WT (blue; 7 slices from 3 animals) and SynTKO (red; 12 slices from 6 animals) recordings. A mixed-effects model revealed no significant difference between genotypes (p = 0.74), but significant differences (p < 0.0001) for the factor time (stimulus). A post-hoc Sidak’s test for multiple comparisons revealed no significant differences for single time points. **c)** Normalized fEPSP amplitudes at the last stimulus of the first stimulation train for individual WT (blue dots; 7 slices from 3 animals) and SynTKO (red dots; 12 slices from 6 animals) recordings. Median values +/- interquartile ranges are shown in black. Ranks were not significantly different (p = 0.1956, Mann-Whitney U test). **d)** Example traces show fEPSP amplitudes of WT (blue) and SynTKO (red) animals in response to the first 10 stimuli of the fourth high-frequency stimulation train. **e)** Normalized averaged fEPSP amplitudes plotted against number of stimuli of the first high-frequency stimulation train for WT (blue; 7 slices from 3 animals) and SynTKO (red; 12 slices from 6 animals) animals. Both the factors time (stimulus), genotype and the interaction of both were significantly different in a mixed-effects model (p < 0.0001 for time, p = 0.02 for the genotype and p = 0.04 for the interaction of genotype and time). A post-hoc Sidak’s test for multiple comparisons revealed no significant differences for single time points. **f)** Normalized fEPSP amplitudes at the last stimulus of the fourth stimulation train for individual WT (blue dots; 7 slices from 3 animals) and SynTKO (red dots; 12 slices from 6 animals) recordings. Median values +/- interquartile ranges are shown in black. Ranks were significantly different (p = 0.004, Mann-Whitney U test).

**Figure 3:**
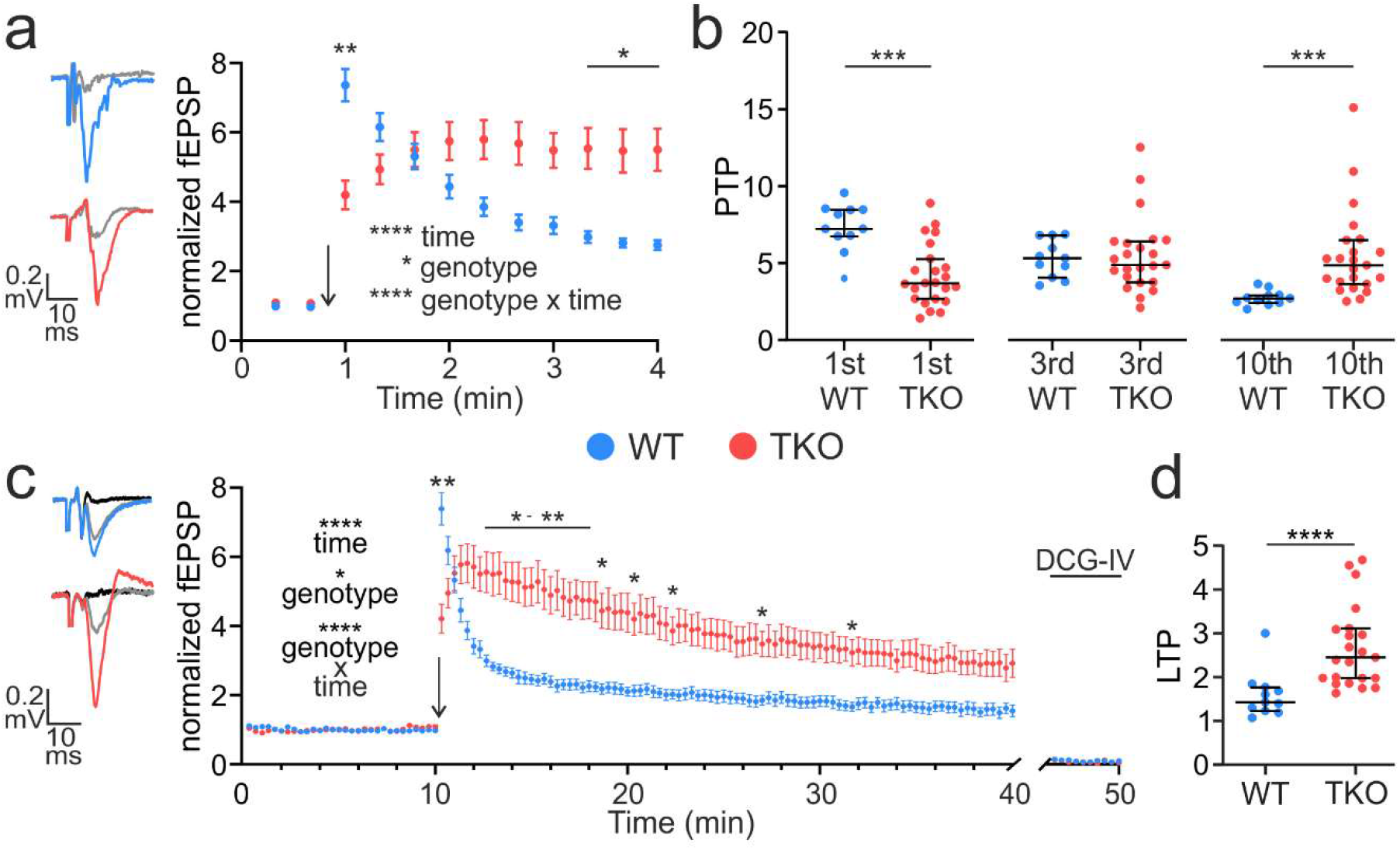
SynTKO mossy fibers display reduced post-tetanic potentiation but increased long-term potentiation. **a)** Post-tetanic potentiation is decreased in SynTKO mossy fibers. **Left:** Example traces of the fEPSP amplitude at the first stimulus after high-frequency stimulation for mossy fibers from WT (blue, top) and TKO (red, bottom) mice compared to averaged baseline fEPSP amplitude (grey). **Right:** Normalized averaged fEPSP amplitudes plotted against time (min) from WT (blue; 11 slices from 4 animals) and SynTKO (red; 23 slices from 9 animals) recordings. This plot is a partial zoom-in from the plot shown in c (minutes 9-13). Mean values +/- SEM are shown. The arrow indicates the time point of high-frequency stimulation (4x 125 pulses @ 25 Hz). Stimulation before and after was at 0.05 Hz. Statistics for data set as reported in c. **b)** Scatter plots for individual fEPSP amplitudes for WT (blue) and SynTKO (red) recordings for the first, third and tenth stimulus after high-frequency stimulation, respectively. Median values +/- interquartile ranges are shown in black. Significance was tested with a Kruskal-Wallis test and a post-hoc Dunn’s correction for multiple comparisons. The Kruskal-Wallis test revealed significant differences between ranks with p < 0.0001. Multiple comparisons revealed significant differences for the first (p = 0.0002) and tenth (p = 0.001) time point. **c)** LTP is increased in SynTKO animals after 30 min. **Left:** Example traces of fEPSP amplitudes 30 min after high-frequency stimulation for mossy fibers from WT (blue, top) and TKO (red, bottom) mice compared to baseline fEPSP amplitude (grey) and response to 1 µM DCG-IV (black). **Right:** Normalized averaged fEPSP amplitudes plotted over time (min) from WT (blue; 11 slices from 4 animals) and SynTKO (red; 23 slices from 9 animals) recordings. Mean values +/- SEM are shown. The arrow indicates the high-frequency stimulation (4x 125 pulses @ 25 Hz). Stimulation frequency before (baseline) and after (LTP recording) was 0.05 Hz. At the end of the recording, 1 µM DCG-IV was washed in to ensure mossy fiber specificity. The last ten fEPSP amplitudes during DCG-IV wash-in are shown at the end of the recording. A mixed-effects model revealed significant differences for the genotype (p = 0.01), time (p < 0.0001) and the interaction of both (p < 0.0001). A post-hoc Sidak’s test for multiple comparisons revealed significant differences for the first sweep after high-frequency stimulation (p = 0.005) and sweeps 38-54 (p: 0.009 – 0.05; ∼13-18 min), as well as for sweeps 56, 61, 67, 81 and 94 (p: 0.03 - 0.04; ∼ minutes 19, 20, 22, 27 and 32). **d)** Dots indicate averaged fEPSP amplitudes from individual WT (blue) and SynTKO (red) recordings. Amplitudes were averaged over the last 10 minutes of the LTP recording; from 20 – 30 min after high-frequency stimulation. Median values +/- interquartile ranges are shown in black. Ranks were significantly different with p < 0.0001 (Mann-Whitney U test).

### R Project for Statistical Computing in RStudio

To account for the multi-level nested structure of the electron microscopy data (Figure 4c,d and Figure 5b), we used a generalized linear mixed model from the gamma family with a log link. We used the glmer function from the R package: lme4 (RRID:SCR_015654) (Bates et al., 2015) to fit the generalized linear mixed models. The different models were compared with an ANOVA. In case of the data underlying Figure 5b we performed a post-hoc test (estimated marginal means with false discovery rate correction) for multiple comparisons. To obtain the marginal means we used the R package: emmeans (RRID:SCR_018734) and compared them in a marginal effects test with false discovery rate correction (Benjamini and Hochberg, 1995).

**Figure 4:**
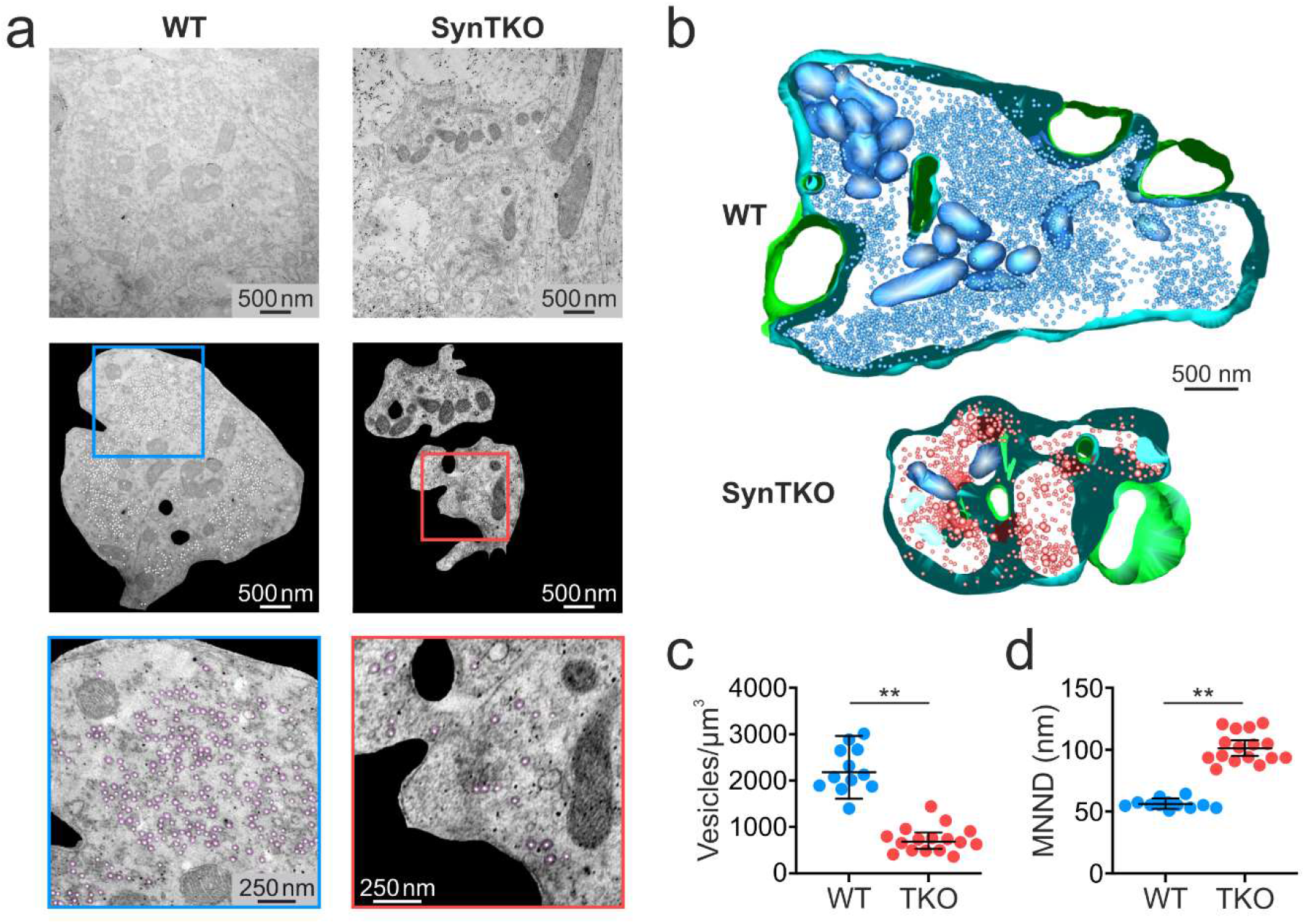
Synaptic vesicles are more dispersed in mossy fiber boutons from SynTKO mice. **a)** In mossy fiber boutons synaptic vesicles are more dispersed and their density is reduced. Example images from transmission electron microscopy (TEM) showing mossy fiber boutons from WT (left) and SynTKO (right) animals. **Top:** Raw TEM images of mossy fiber boutons in stratum lucidum. **Middle:** An automated tool (Imbrosci et al., 2022) was used to detect vesicles. Mossy fiber boutons were extracted from the raw image and the center of detected vesicles is marked with a white dot. Blue and red boxes show the region for the zoom-ins in WT and SynTKO, respectively. **Bottom:** Zoom-ins, as marked in the middle pictures. Vesicles detected by the algorithm are shown in purple. **b)** Partial 3D reconstruction of hippocampal mossy fiber boutons from a WT (top) and a SynTKO animal (bottom) for visualization purposes only. Vesicles are shown in blue and red respectively, the presynaptic mossy fiber membrane is shown in light blue and postsynaptic spines are shown in green. **c)** The number of synaptic vesicles per µm^3^ is reduced in SynTKO animals. Dots represent the number of vesicles in individual mossy fiber boutons from 2 WT (blue; 12 boutons) and 3 SynTKO (red; 16 boutons) animals. Mean values and the borders of the 95% confidence intervals are shown in black. A generalized linear mixed model (gamma family with log link) revealed significant differences between genotypes with p = 0.006. **d)** The mean nearest neighbor distance (MNND) is increased between synaptic vesicles in SynTKO compared to WT boutons. Scatter plot shows average MNND (nm) for individual mossy fiber boutons from 2 WT (blue; 12 boutons) and 3 SynTKO (red; 16 boutons) animals. Genotypes were significantly different in a generalized linear mixed model (gamma family with log link) with p = 0.0015. Mean values are shown in black with the borders of the 95% confidence intervals.

**Figure 5:**
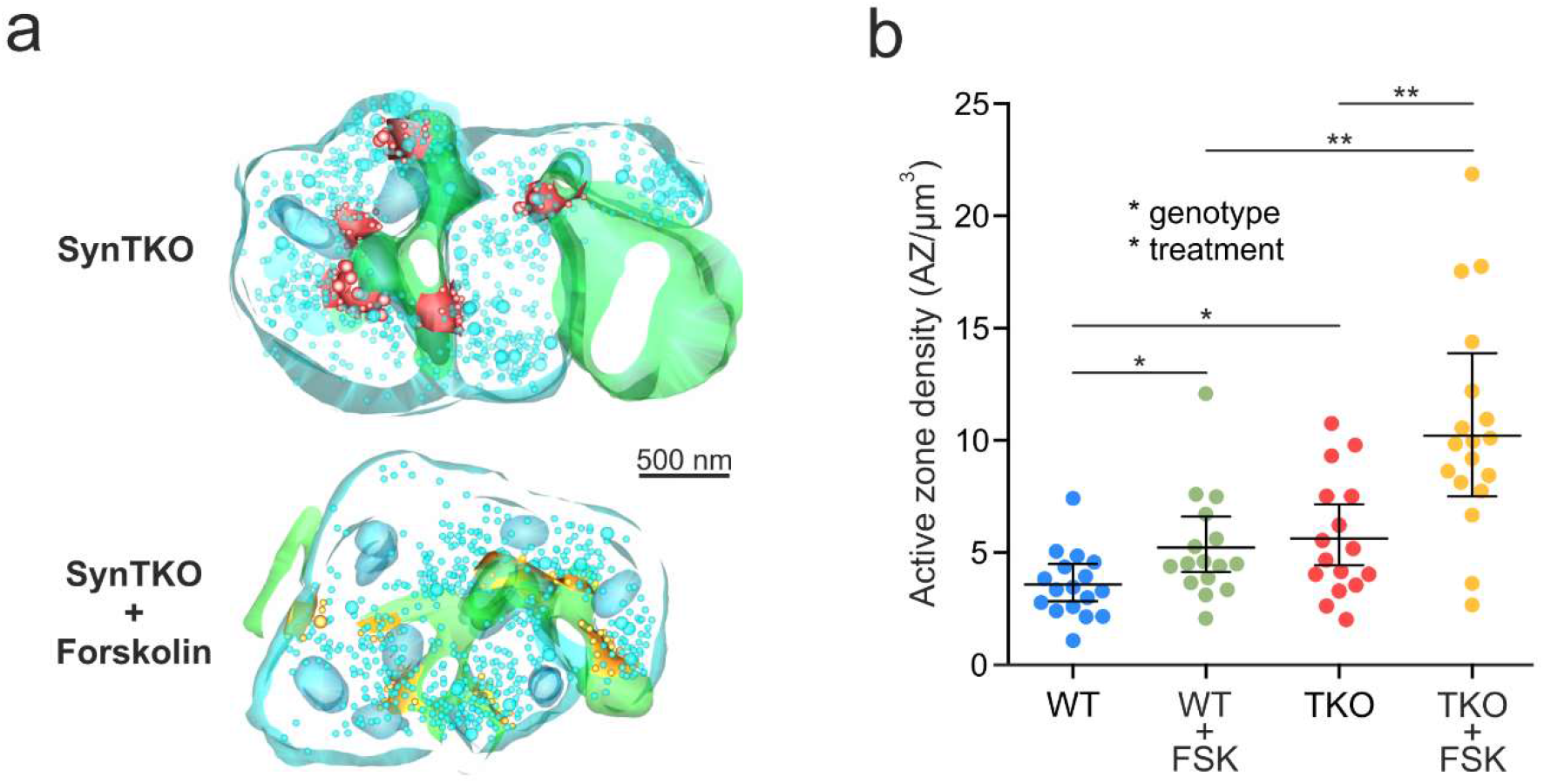
Increased active zone density in mossy fiber boutons of SynTKO mice. **a)** Example partial 3D reconstructions of mossy fiber boutons from untreated (top) and forskolin-treated (bottom) SynTKO mice. Active zones with docked vesicles are shown in red (SynTKO) and yellow (SynTKO + forskolin), respectively. Synaptic vesicles, mitochondria and presynaptic membrane are shown in light blue, postsynaptic membrane is shown in green. **b)** Number of active zones per µm^3^ plotted for individual mossy fiber boutons from untreated WT (blue dots; 17 boutons from 3 animals) and untreated SynTKO slices (red dots; 16 boutons from 3 animals) as well as for forskolin-treated WT (green dots; 16 boutons from 3 animals) and forskolin-treated SynTKO slices (yellow dots; 18 boutons from 3 animals). Mean values are shown in black with upper and lower borders of the 95% confidence interval. A generalized linear mixed model (gamma family with log link) revealed significant differences for genotype (p = 0.01) and forskolin treatment (p = 0.013). A post-hoc test (estimated marginal means with false discovery rate correction) revealed significant differences between WT and SynTKO (p = 0.0125), WT and WT + forskolin (p = 0.0134), SynTKO and SynTKO + forskolin (p = 0.0018) and WT + forskolin and SynTKO + forskolin (p = 0.0018), but no significant difference between WT + forskolin and SynTKO (p = 0.658).

## Results

### Mossy fibers of SynTKO animals are more excitable

Despite the sparse connectivity (Amaral et al., 1990) and the low baseline activity of granule cells (Jung and McNaughton, 1993), a single mossy fiber *bouton* is able to trigger the discharge of its postsynaptic partner (Henze et al., 2002; Vyleta et al., 2016). Mossy fiber activity is not only important for pattern separation in the healthy brain (Rolls, 2018), but also for propagation of seizures in the epileptic brain (Nadler, 2003). Since SynTKO animals display high network excitability and develop epileptic seizures at the age of two months (Gitler et al., 2004; Fassio et al., 2011), we tested the excitability in mossy fibers by measuring the input-output relationship. We performed experiments in presymptomatic (4-6 weeks old) and symptomatic (17-19 weeks old) animals, after the onset of epileptic seizures. This design was aimed at differentiating changes in synaptic transmission that could lead to or result from epilepsy in SynTKO animals.

We conducted field recordings in acute slices from SynTKO and wildtype (WT) age-matched controls and recorded the input-output relationship as a measure of synaptic strength. We recorded from the *stratum lucidum* of area CA3 while stimulating close to the granule cell layer in the hilus (Figure 1a). We found that SynTKO were significantly more excitable than WT animals: the input-output relation was increased in both 4-6 weeks old (Figure 1b) and 17-19 weeks old animals (Figure 1-1b). With the same amount of stimulated fibers (size of the presynaptic fiber volley), the excitatory postsynaptic field potential (fEPSP) amplitudes were larger. The slopes of the simple linear regression fits of fiber volley versus fEPSP amplitudes were significantly different with p < 0.0001 between control and SynTKO data for both age groups. For presymptomatic recordings the slopes for the simple linear regression with 95% confidence intervals were 0.106 [0.018 – 0.194] for WT and 2.743 [2.159 – 3.326] for SynTKO and for symptomatic recordings 0.795 [0.59 – 0.99] for WT and 2.292 [1.806 – 2.777] for SynTKO.

The possible reasons underlying this increased excitability are manifold. Since the hippocampal morphology was described to be similar between WT and SynTKO animals (Gitler et al., 2004), we assume that the number of excitable fibers is comparable. To check for a possible change in release probability, we measured the paired-pulse ratio (PPR) with an inter-stimulus interval of 50 ms. Under our experimental conditions the PPR was not significantly different between presymptomatic SynTKO and WT animals (Figure 1-2a). In WT recordings the median PPR was 2.962 [2.510; 3.960], while in recordings from SynTKO it was 3.745 [2.422; 5.570] with a p-value > 0.2 (Mann-Whitney *U* test). However, we saw a trend for an increased PPR that became clearer with a shortening of the inter-stimulus interval to 40 ms (see first two fEPSPs in Figure 2a,b). Here, in WT recordings, the median PPR was 3.833 [2.654; 4.026], while it was 4.436 [3.485; 6.094] for SynTKO (Figure 1-2b). The p-value was 0.14 (Mann-Whitney *U* test). Finally, in symptomatic SynTKO animals, the PPR was significantly increased compared to WT animals with p = 0.03 (Mann-Whitney *U* test) with a median of 2.906 [2.549; 3.499] for WT and 3.435 [2.964; 4.944] for SynTKO animals (Figure 1-2c). A change in PPR is suggestive of a change in release probability (Dobrunz and Stevens, 1997); however, this might be taken with caution as many other factors affect this measure (Hanse and Gustafsson, 2001; Sun et al., 2005; Neher and Brose, 2018; Glasgow et al., 2019).

### Reduced frequency facilitation in mossy fibers of SynTKO animals

Mossy fiber *boutons* are very powerful synapses when it comes to presynaptic plasticity. They are able to facilitate dramatically, even at moderate frequencies (Salin et al., 1996). This phenotype, together with large pools of synaptic vesicles (Hallermann et al., 2003; Rollenhagen et al., 2007), makes them an excellent system for studying the influence of synapsins on presynaptic plasticity. In previous work, it has been reported that frequency facilitation is impaired at mossy fibers from SynDKO animals after stimulation with a moderate frequency of 2 Hz (Owe et al., 2009). The authors suggested that the remaining synapsin isoform – SynIII – causes impaired facilitation since it was localized in the RRP of mossy fiber *boutons*. Additionally, neurons from SynIII KO animals show less synaptic depression than WT neurons in primary hippocampal cultures (Feng et al., 2002). Here, we intended to test if complete deletion of synapsins, including SynIII, would rescue frequency facilitation at the hippocampal mossy fiber *bouton*.

When stimulated with a train of 20 pulses at a frequency of 1 Hz, we saw less facilitation in mossy fibers from presymptomatic SynTKO compared to WT animals. This finding is comparable to the aforementioned experiments in SynDKO animals (Owe et al., 2009) and cell culture experiments of SynTKO animals (Gitler et al., 2004). The rise in the field excitatory postsynaptic potential (fEPSP) amplitudes was similar in WT and SynTKO during the first 10 stimuli, but in SynTKO animals we observed an earlier saturation of amplitudes. In SynTKO the amplitudes reached a plateau after 15 stimuli, whereas in WT animals the amplitudes increased until the end of the 1 Hz stimulation (Figure 1c). When comparing the plots with a mixed-effects model, we found significant differences for the factor time, as well as for the interaction between time and genotype (p < 0.0001). The post-hoc Sidak’s test for multiple comparisons revealed no significant differences for single time points (p > 0.05). When comparing only the amplitudes in response to the last 1 Hz stimulus, we found that the median facilitation was 6.909 [5.589; 9.093] for WT animals, while the increase was only 5.110 [3.760; 7.080] compared to the baseline for SynTKO animals (median value [25% quartile; 75% quartile]). Ranks were different with p = 0.0029 (Mann-Whitney *U* test (Figure 1d)).

This result suggests that SynIII is not the primary reason for the reduced facilitation in mossy fiber *boutons*. However, Owe and coworkers used three to six months old SynDKO mice. Since all synapsin knockout animals lacking SynI and/or SynII develop seizures beginning at the age of two months (Fassio et al., 2011), this pathology could potentially lead to secondary differences in plasticity. We investigated short-term plasticity also in 17-19 weeks old symptomatic mice – matching the age range from Owe et al. – and observed an even more pronounced effect on frequency facilitation (Figure 1-1c,d): the facilitation in WT animals reached 7.019 [5.574; 8.440] while the increase in SynTKO recordings was only 4.414 [4.036; 5.330] compared to baseline (median [25% quartile; 75% quartile]). Ranks were significantly different with p = 0.0009 (Mann-Whitney *U* test). Thus, the additional knockout of SynIII did not lead to a rescue of facilitation, neither in presymptomatic nor in symptomatic animals. Hence, our data do not support the hypothesis that SynIII acts as a brake on frequency facilitation in hippocampal mossy fiber *boutons*.

In summary, we see a decrease in frequency facilitation, but an increase in excitability in the absence of synapsins. These results indicate that, (1) SynIII is not causing the reduced facilitation and (2) that before the onset of epileptic seizures, excitability and short-term plasticity mechanisms are already altered.

### High-frequency stimulation leads to early vesicle exhaustion in SynTKO animals

Since stimulation with a moderate frequency led to a decrease in facilitation in presymptomatic SynTKO animals (Figure 1c,d), we wanted to investigate the response to a longer stimulation with a higher frequency. We applied four trains of 125 pulses at 25 Hz using the same recording paradigm as before (Figure 1a). While the course of the amplitudes was very similar in the first high-frequency train for both genotypes (Figure 2a,b,c), changes manifested over time. Differences between WT and SynTKO animals became distinguishable in the fourth stimulation train (Figure 2d,e,f) with smaller amplitudes throughout the whole train in SynTKO animals.

When tested with a mixed-effects model, we detected significant differences for the factor time (stimuli) for both the first and the fourth stimulation train (p < 0.0001). The factor genotype and the interaction of genotype and time were only significant for the fourth stimulation train (p < 0.05). A post-hoc Sidak’s test for multiple comparisons revealed no significant differences for single time points for either of the stimulation trains. We also compared the amplitudes for the last stimulus of the stimulation trains between genotypes. For the first stimulation train, the median normalized fEPSP amplitude was 1.848 [1.243; 2.467] for WT, while it was 0.7514 [0.4997; 1.650] for SynTKO animals. Ranks were not significantly different (p = 0.1956; Mann-Whitney *U* test) (Figure 2c). However, when comparing the amplitudes of the last stimulus of the fourth stimulation train, we found a significant difference (p = 0.0037, Mann-Whitney *U* test) between WT and SynTKO (Figure 2f). The median normalized fEPSP amplitude was 1.381 [0.6876; 2.293] for WT and 0.3714 [0.04892; 0.7202] for SynTKO animals. This stronger exhaustion during intense stimulation was already described before in other synapsin knockout animals (Rosahl et al., 1995; Farisello et al., 2013) and probably reflects the missing reserve pool, which would normally replenish the RRP under such high activity (Vasileva et al., 2012).

High-frequency stimulation can lead to a loss of fibers during the course of stimulation. To check if the smaller fEPSP amplitudes in the last stimulation train of SynTKO recordings is due to an increased fiber loss, we measured a subset of the presynaptic fiber volleys (PFV) during the first and fourth stimulation train for both genotypes, respectively. While PFVs of SynTKO animals were in general smaller than the ones from WT animals with comparable fEPSP amplitudes (Figure 1b), there was no relative difference in PFV sizes of the two genotypes between first and last stimulation train (Figure 2-1). Thus, we conclude that the relative loss of fibers is similar for both genotypes and does not explain the more drastic decrease in fEPSP size for SynTKO animals.

Here, our data indicate that deletion of all synapsins disturbs the vesicle organization in synaptic terminals in a way that leads to impaired replenishment. This is especially relevant for synapses like mossy fiber *boutons*, which have large vesicle pools (Hallermann et al., 2003; Rollenhagen et al., 2007).

### Post-tetanic potentiation is changed in SynTKO animals

After intense stimulation of mossy fibers, another form of short-term plasticity occurs: post-tetanic potentiation (PTP) (Griffith, 1990), which was proposed to underlie short-term memory (Vandael et al., 2020). Measuring PTP after four trains of high-frequency stimulation revealed differences between WT and SynTKO animals: while in WT recordings the median potentiation was 7.234 [6.752; 8.478] fold compared to baseline and decreased over time, in SynTKO recordings, we initially measured an amplitude which was only 3.702 [2.683; 5.280] times larger than baseline (significantly different in a Kruskal-Wallis test with post-hoc Dunn’s test for multiple comparisons; p = 0.0002), but increased over time. After one minute, the amplitudes of WT and SynTKO recordings were comparable (Figure 3a,b; WT: 5.323 [4.070; 6.812]; SynTKO: 4.887 [3.769; 6.419]; p > 0.99), followed by a further increase in the SynTKO amplitudes over WT amplitudes. One minute after stimulation, the amplitudes of the SynTKO animals remained on a plateau while the amplitudes in the WT animals decreased further (Figure 3a,b), leading to median amplitudes of 2.717 [2.432; 2.922] for WT and 4.865 [3.635; 6.497] for SynTKO animals approximately three minutes after high-frequency stimulation (significantly different with p = 0.001). These findings point to different underlying mechanisms: one leading to the impairment of PTP right after high-frequency stimulation and another one leading to increased amplitudes after some recovery time and upon low frequency stimulation of 0.05 Hz. To understand this observation further, we continued recording for half an hour, which corresponds to early long-term potentiation (LTP).

### Long-term potentiation is enhanced in SynTKO animals

Mossy fiber *boutons* express a presynaptic form of LTP, which is PKA-dependent (Weisskopf et al., 1994). In recordings from SynDKO animals mossy fiber LTP was unchanged compared to WT animals (Spillane et al., 1995). It is tempting to speculate, though, that SynIII might be the phosphorylation target of PKA in the context of LTP, since a PKA phosphorylation site is present in domain A (Piccini et al., 2015), which is conserved among all synapsins, and SynIII expression is maintained in adult mossy fiber *boutons* (Pieribone et al., 2002). An additional knockout of SynIII could therefore lead to a block of LTP. However, when recording long-term potentiation (Figure 3c) we measured a median potentiation of 1.43 [1.23; 1.77] in WT animals 20 to 30 minutes after the high-frequency stimulation, while SynTKO animals showed a larger median potentiation of 2.45 [1.98; 3.12] compared to baseline (Figure 3d). Ranks differed significantly with p < 0.0001 (Mann-Whitney *U* test). The time course of LTP was tested in a mixed-effects model. The factors genotype, time and the interaction of both differed significantly (p = 0.013; p < 0.0001; p < 0.0001, respectively). A post-hoc Sidak’s test for multiple comparisons revealed significant differences for single time points as well (Figure 3c). We included all measurements that fulfilled the specificity criterion, which was tested by the application of the metabotropic glutamate receptor group II agonist (2*S*,1’*R*,2’*R*,3’*R*)-2-(2,3-dicarboxycyclopropyl)glycine (DCG-IV) (Kamiya et al., 1996) (last ten sweeps are shown in Figure 3c).

In summary, the absence of all synapsin isoforms in mossy fiber synapses leads to a reduced early PTP, an altered time-course of PTP/LTP and an increased long-lasting potentiation. Such changes in LTP have not been described before in other synapsin knockout models, suggesting that it is an effect of SynIII deletion and specifically relevant for the mossy fiber *bouton,* where LTP occurs presynaptically (Zalutsky and Nicoll, 1990). Since it has been shown that ultrastructural changes underlie potentiation at hippocampal mossy fibers (Orlando et al., 2021) we next sought to investigate the ultrastructure of mossy fiber *boutons* in SynTKO animals.

### Synaptic vesicles are more dispersed in SynTKO animals

So far, vesicle distributions at the hippocampal mossy fiber *bouton* have only been described for either SynDKO animals or SynIII KO animals (Feng et al., 2002; Owe et al., 2009). Here, we wanted to test whether the knockout of all three synapsins would lead to additional changes in vesicle organization at the hippocampal mossy fiber *bouton*. Using transmission electron microscopy (TEM), we identified individual mossy fiber *boutons* from three presymptomatic SynTKO and two age-matched WT mice. We imaged serial sections from 16 SynTKO and 12 WT mossy fiber *boutons*. For each 2D projection, we measured the vesicle number and the mean nearest neighbor distance (MNND) using an automated tool (Imbrosci et al., 2022) (Figure 4a). For both data sets, we used a generalized linear mixed model (gamma family with log link) to predict either the vesicle density or MNND. When comparing synaptic vesicles of WT and SynTKO *boutons*, the mean density was strongly reduced in *boutons* from SynTKO animals (687 [532; 886] vesicles/µm^3^ compared to 2186 [1612; 2965] vesicles/µm^3^ in WT) (Figure 4b,c). The genotypes were significantly different with p = 0.006. Consequently, we also saw an increase in the MNND of vesicles (Figure 4b,d): the average MNND was 56.3 [52.3; 60.6] nm for WT and 101.2 [95.1; 107.8] nm for SynTKO *boutons*. Groups were significantly different with p = 0.0015. The reduced density of distal vesicles implies a reduced reserve pool. Since this observation resembles the results seen in mossy fiber *boutons* of SynDKO animals (Owe et al., 2009), our data indicate that the additional knockout of SynIII does not add on effects on the organization of the distal pool. This conclusion is also in line with unchanged synaptic vesicle densities in mossy fiber *boutons* of SynIII KO mice (Feng et al., 2002).

### Active zone density is highest in chemically potentiated mossy fiber *boutons* from SynTKO animals

Since we saw an increase in LTP in SynTKO animals (Figure 3c,d), we wanted to understand if structural changes would occur in potentiated mossy fiber *boutons* from SynTKO animals. We performed TEM in hippocampal slices from both young WT and SynTKO animals in either potentiated or control conditions. Potentiation was chemically induced via incubation with the adenylyl cyclase-activator forskolin before fixation of the samples. Forskolin has similar effects on mossy fibers as high-frequency electrical stimulation (Weisskopf et al., 1994; Spillane et al., 1995). The active zone density was analyzed in partial 3D reconstructions of mossy fiber *boutons* from three animals per group and replicates of 17 (WT), 16 (WT + forskolin), 16 (SynTKO) and 18 (SynTKO + forskolin) *boutons*, respectively. We fitted a generalized linear mixed model (gamma family with a log link) to predict active zone density with genotype and forskolin treatment, which included the individual animals as random effects. We found significant differences for genotype (p = 0.01) and forskolin treatment (p = 0.013). Specific pairs were compared by testing estimated marginal means with adjustment for false discovery rate (Benjamini and Hochberg, 1995).

We observed an increase in the active zone density in WT animals when treated with forskolin, as described before (Orlando et al., 2021). Untreated *boutons* from SynTKO animals had a similar mean density of active zones as forskolin-treated *boutons* from WT animals (5.63 [4.43; 7.14] active zones/µm^3^ for untreated SynTKO *boutons*; 5.22 [4.13; 6.60] active zones/µm^3^ for forskolin-treated WT *boutons,* p = 0.658). This indicates that, from a structural point of view, SynTKO animals could be in a similar state as potentiated WT *boutons*. Treatment with forskolin led to a further increase in the active zone density in mossy fiber *boutons* from SynTKO animals (10.20 [7.50; 13.88] active zones/µm^3^) (Figure 5a) and led to significant differences when compared to untreated SynTKO *boutons* (p = 0.0018) as well as treated WT *boutons* (p = 0.0018) (Figure 5b). Taken together, we might see a structural strengthening in *boutons* from SynTKO animals treated with forskolin, which could explain the increase in long-term potentiation (Figure 3c,d).

## Discussion

Here, we demonstrate that synapsin-dependent vesicle organization plays a crucial role in various forms of presynaptic plasticity at hippocampal mossy fiber *boutons*. The removal of all synapsin isoforms leads to impaired PTP, indicating a potential role of synapsins in short-term memory. Active zone density was increased in mossy fiber *boutons* of SynTKO animals, indicating a preset potentiated state. This morphological phenotype might underlie the increased LTP we observed in SynTKO animals. Together, our results indicate that all synapsin isoforms, including SynIII, play a role in the modulation of mossy fiber-specific presynaptic plasticity.

In SynTKO mice, we found increased excitability, measured by a change in the input-output relation of local fEPSPs (Figure 1, Figure 1-1). A likely explanation is based on the finding that synapsins play different roles in excitatory versus inhibitory neurons (Song and Augustine, 2015). Deletion or mutation of SynI, SynIII or all synapsins leads to impaired basal transmission of inhibitory, but not excitatory cultured neurons (Terada et al., 1999; Feng et al., 2002; Gitler et al., 2004; Baldelli et al., 2007) and loss of SynII impairs tonic inhibition in hippocampal slices (Medrihan et al., 2013, 2015). Mossy fibers activate at least four times more inhibitory neurons than pyramidal cells in CA3 (Acsády et al., 1998), regulating CA3 excitability via feedforward inhibition (Acsády and Káli, 2007; Torborg et al., 2010). Reduced feedforward inhibition might thus explain the increased excitability. Indeed, the input-output relation is increased in Schaffer collaterals from SynTKO animals, while it is reduced in inhibitory fibers from CA1 (Farisello et al., 2013).

During trains of activity, mossy fiber *boutons* facilitate reliably (Salin et al., 1996; Toth et al., 2000), which is thought to be important for information transfer (Henze et al., 2002; Mori et al., 2004). In mossy fibers from SynDKO animals, frequency facilitation is reduced (Owe et al., 2009). There, the authors suggested that the remaining SynIII may act as a brake on facilitation, because (1) SynIII is associated specifically with the RRP in mossy fiber *boutons* (Owe et al., 2009) and (2) synaptic depression is reduced in SynIII KO cultures (Feng et al., 2002). However, in animals lacking all synapsins, including SynIII, we still observed reduced frequency facilitation (Figure 1, Figure 1-1), rejecting the hypothesis from Owe and colleagues. Frequency facilitation is most likely calcium-dependent and involves increased neurotransmitter release (Chamberland et al., 2017; Jackman and Regehr, 2017). Hence, potential reasons for reduced facilitation are diverse and include enhanced basal release probability, depletion of the RRP and saturation of postsynaptic receptors (Neher and Sakaba, 2008).

High-frequency stimulation usually results in a biphasic depression, attributed to the depletion of the RRP (Zucker and Regehr, 2002) and slow replenishment from the reserve pool (Wesseling and Lo, 2002). We observed frequency-dependent depression for both genotypes when stimulating at 25 Hz (Figure 2), but stronger depression in SynTKO animals, recapitulating previous results (Gitler et al., 2004). At the calyx of Held, a reduced reserve pool and slower replenishment accounted for faster depression in SynTKO animals (Vasileva et al., 2012). Indeed, in mossy fiber *boutons* of SynTKO animals, vesicles in the distal pool were reduced in density and more dispersed (Figure 4), likely explaining faster depression. Impaired distal pools were described before for mossy fiber *boutons* (Takei et al., 1995; Owe et al., 2009) and in neuronal cultures from SynTKO mice (Gitler et al., 2004; Siksou et al., 2007). Hence, at mossy fibers, the additional knockout of SynIII recapitulates previously described phenotypes – in line with unchanged reserve pools in SynIII KO mossy fiber *boutons* (Feng et al., 2002) and a role of SynIIa in vesicle replenishment (Gitler et al., 2008). In general, vesicle de-clustering and reduced vesicle density likely have diverse effects on the release cycle (Bykhovskaia, 2011), possibly also supporting increased excitability and reduced frequency facilitation.

PTP has recently been suggested to underlie short-term memory. During mossy fiber PTP a “pool engram” is formed, i.e. the number of docked vesicles at active zones increases (Vandael et al., 2020). This engram formation depends on the refilling rate of vesicles and could thus be mediated by synapsins. Our data support this hypothesis: the complete loss of synapsins impairs mossy fiber PTP (Figure 3). Reduced PTP was observed before in synapsin KO models, with diversity regarding synapsin isoform and synapse type. PTP is reduced in (1) cultured hippocampal neurons of SynI KO and SynTKO animals (Valente et al., 2012; Cheng et al., 2018), (2) at Schaffer collaterals of SynII KO, SynDKO and SynTKO animals (Rosahl et al., 1995; Farisello et al., 2013) and (3) at corticothalamic synapses of SynI, but not SynII, KO animals (Nikolaev and Heggelund, 2015). However, no change in PTP was reported for mossy fibers of SynDKO animals (Spillane et al., 1995). We therefore speculate that the lack of SynIII in SynTKO mice might cause the additional PTP phenotype that we observe in mossy fibers. In cell culture, PTP measured via miniature excitatory postsynaptic currents could only be rescued by the SynIIIa isoform (Cheng et al., 2018), supporting this notion.

While the initial drop in PTP could be explained by impaired vesicle replenishment (Vasileva et al., 2012), we also observed a second, increased PTP phase (Figure 3a,b). Alongside the RRP, also release probability and quantal size are increased during mossy fiber PTP (Vandael et al., 2020). Both could be elevated by default in SynTKO animals and elevate PTP in the second phase. Interestingly, we detected an increase in active zone density in SynTKO *boutons* (Figure 5), which most likely reflects a change in the number of release sites. Hence, after overcoming the initial drop in PTP, other mechanisms could be untamed in SynTKO mossy fiber *boutons*, leading to enhanced PTP in a later phase.

We discovered previously that active zone density is increased in potentiated mossy fiber *boutons* (Orlando et al., 2021). Therefore, the increased density in SynTKO animals could indicate a preset potentiated state due to homeostatic adaptation, similar to mechanisms in the calyx of Held of SynTKO animals (Vasileva et al., 2012). The active zone density was further increased when chemically potentiating SynTKO mossy fibers with forskolin, leading to significantly higher densities than in forskolin-treated WT *boutons* and untreated SynTKO *boutons* (Figure 5). Forskolin could induce these structural changes via SynIII phosphorylation by PKA, similar to developmental processes (Piccini et al., 2015). Elevated active zone densities might also explain the increased LTP we observe in SynTKO animals (Figure 3). Mossy fiber LTP was analyzed before in SynI KO (Takei et al., 1995) and SynDKO mice (Spillane et al., 1995), but was found to be unchanged. Thus, we speculate that the increase in LTP is a likely consequence of the additional knockout of SynIII.

It is unknown which mechanisms are shared between mossy fiber PTP and LTP and when one results in the other. SynIII might have a specific function in both processes, preventing excess release and balancing potentiation. Recent literature suggests that (1) diversity in STP depends on priming and fusion steps (Lin et al., 2022) and (2) increased fusion competence might underlie mossy fiber LTP, possibly mediated by Munc13-1 (Lipstein et al., 2021; Fukaya et al., 2023a; Papantoniou et al., 2023). Does SynIII play a role in the insertion of new active zones, vesicle docking, priming and/or fusion? Such a role would most likely be intermingled with SynIIIs role in neurogenesis (Kao et al., 2008). Future work in SynIII KO models will allow us to answer this question.

Here, we investigated plasticity at a glutamatergic synapse expressing SynIII in adulthood. We used SynTKO instead of SynIII KO animals to exclude compensatory effects via remaining synapsin isoforms. However, this approach also limits our ability to draw precise conclusions on SynIII-specific effects. By combining physiological recordings – well-suited to record mossy fiber transmission (Breustedt et al., 2010) – and ultrastructural studies, our experiments shed light on synapsin-dependent plasticity from different angles. To exclude possible indirect estrogen-effects on mossy fiber plasticity (Harte-Hargrove et al., 2013), we used male mice only, limiting the generalizability. Future studies should include female animals. Finally, although chemical mossy fiber potentiation is widely used, it is still unclear if it shares the same mechanisms as electrical induction (Shahoha et al., 2022; Fukaya et al., 2023b).

Our work revealed that the complete loss of synapsins leads to disruption of presynaptic plasticity at hippocampal mossy fibers. Facilitation and PTP are reduced, likely due to an impaired vesicle replenishment. However, LTP is increased, in concert with an elevated active zone density. We speculate that the loss of SynIII supports these physiological and ultrastructural changes. Our work contributes to a better understanding of mossy fiber presynaptic plasticity and, consequently, to a better understanding of synapsins’ roles in learning and memory. Further work is needed to dissect the precise roles of the various synapsin isoforms both in hippocampal mossy fiber *boutons* and in other synapses, especially those expressing SynIII in adult stages (Pieribone et al., 2002).

## Supplementary figures

**Figure 1-1: Increased excitability, but reduced facilitation, at mossy fibers of symptomatic SynTKO. a)** Field recording setup for an acute brain slice. The stimulation electrode was placed in the hilus close to the granule cell layer of the dentate gyrus, the recording electrode was placed in the stratum lucidum of area CA3 of the hippocampus where mossy fibers terminate. **b)** Excitability was increased in recordings from SynTKO mice (red; 18 slices from 5 animals) compared to WT mice (blue; 17 slices from 4 animals). Pooled fEPSP amplitudes (mV) were plotted against pooled presynaptic fiber volley (PFV) amplitudes (mV) and fitted with a simple linear regression. The slopes of the linear regressions were significantly different (p < 0.0001, tested with a two-tailed ANCOVA). 95% confidence bands are shown as dotted lines around the fit. **Inset:** Example traces from WT (blue) and SynTKO (red) animals with similar PFV amplitudes. Note the difference in the corresponding fEPSP amplitude. **c)** Frequency facilitation was reduced in SynTKO (red) compared to WT (blue) animals. Averaged normalized fEPSP amplitudes +/- SEM from all WT (blue; 17 slices from 4 animals) and SynTKO (red; 19 slices from 5 animals) recordings plotted against the number of stimuli. Stimuli 1-10 were given with a frequency of 0.05 Hz, stimuli 11-30 with 1 Hz and stimuli 31-41 with a frequency of 0.05 Hz again. Both time and the interaction between genotype and time were significantly different in a mixed-effects model (p < 0.0001). Post-hoc Sidak’s test for multiple comparisons revealed significant differences (p < 0.05) for two time points (indicated with *). **d)** Facilitation was reduced in SynTKO compared to WT animals after moderate frequency stimulation. **Left:** fEPSP amplitudes at the 20^th^ stimulus at 1 Hz for individual WT (blue dots; 17 slices from 4 animals) and SynTKO (red dots; 19 slices from 5 animals) recordings. Median values +/- interquartile ranges are shown in black. Facilitation was significantly different (p = 0.0009, tested with Mann-Whitney U test). **Right:** Example fEPSP amplitudes from WT (blue) and SynTKO (red) animals at the 20^th^ 1 Hz stimulus. Respective baseline fEPSP amplitudes are shown in grey.

**Figure 1-2: Paired-pulse ratio not significantly changed in presymptomatic SynTKO mice. a)** Paired-pulse ratio for presymptomatic SynTKO and age-matched control animals. **Top:** Example traces for a paired-pulse from WT (blue) and SynTKO (red) recordings, respectively. **Bottom:** Dots represent paired-pulse ratios from individual recordings from WT (blue dots, 26 slices from 10 animals) and SynTKO (red dots, 34 slices from 12 animals) slices, calculated as the ratio of second to first fEPSP amplitude. The inter-stimulus interval (ISI) was 50 ms. Median values +/- interquartile ranges are depicted in black. Ranks were compared in a Mann-Whitney U test and were not significantly different (p = 0.226). **b)** Paired-pulse ratio for presymptomatic SynTKO (red) animals and age-matched controls (blue) with a shorter ISI. Dots represent individual paired-pulse ratio of the first two stimuli from the 25 Hz stimulation train (Figure 2a,b) with an ISI of 40 ms, for WT (blue dots, 7 slices from 3 animals) and SynTKO (red dots, 12 slices from 6 animals) recordings. Median values +/- interquartile ranges are depicted in black. Ranks were tested with a Mann-Whitney U test and were not significantly different (p = 0.142). **c)** Paired-pulse ratio for symptomatic SynTKO and age-matched control animals. **Top:** Example traces for a paired-pulse from WT (dark blue) and SynTKO (dark red) recordings, respectively. **Bottom:** Dots represent paired-pulse ratios from individual recordings from WT (dark blue dots, 17 slices from 4 animals) and SynTKO (dark red dots, 19 slices from 5 animals) slices, calculated as the ratio of second to first fEPSP amplitude. The inter-stimulus interval (ISI) was 50 ms. Median values +/- interquartile ranges are depicted in black. Ranks were compared in a Mann-Whitney U test and were significantly different (p = 0.0325).

**Figure 2-1: The loss of fibers during high-frequency stimulation is not substantial and similar for SynTKO and WT mice. a)** Exemplary traces from high-frequency stimulation trains for WT (blue) and SynTKO (red) animals. The 10^th^ PFV and fEPSP from the first and fourth stimulation train are depicted, respectively. Dotted lines indicate the peaks of the PFV. Note that although the PFV is smaller for SynTKO (due to technical reasons in response to the high excitability), the size is relatively consistent throughout the trains. **b)** Averaged PFV (mV) taken from pulses 10-15 from the first and fourth stimulation train, respectively, for recordings from WT (blue; 7 slices from 3 animals) and SynTKO (red; 12 slices from 6 animals) slices. Average values from the same recording are connected. Median values +/- interquartile ranges are depicted in black. Ranks between first and fourth stimulation train were not significantly different for neither WT (p = 0.16) nor SynTKO (p = 0.08) recordings, compared with a Wilcoxon test. **c)** The relative loss of fibers was similar for WT and SynTKO recordings. Averaged ratios between 4^th^ and 1^st^ train PFV sizes are depicted for both WT (blue; 7 slices from 3 animals) and SynTKO (red; 12 slices from 6 animals) animals. Median values +/- interquartile ranges are depicted in black. Ranks were not significantly different (p = 0.45) in a Mann-Whitney U test.

## Acknowledgements

This study was supported by the Deutsche Forschungsgemeinschaft (DFG, German Research Foundation), project 184695641 – SFB 958 (to D.S.), project 327654276 – SFB 1315 (to D.S.), project 415914819 – FOR 3004 (to D.S.), project 431572356 (to D.S.), under Germany’s Excellence Strategy EXC-2049-390688087 (NeuroCure; to D.S. and M.O), project 503954250 (to M.O.). It was also supported by the European Research Council (ERC) under the European Union’s Horizon 2020 research and innovation program (BrainPlay Grant agreement No. 810580; to D.S.) and by the Federal Ministry of Education and Research (BMBF, SmartAge – project 01GQ1420B; to D.S.). D.M. is supported by the start-up funds from DZNE, the grants from the German Research Foundation (SFB 1286/B10 and MI 2104), and the Human Frontiers Science Organization (RGEC32/2023). The funders had no role in study design, data collection and analysis, decision to publish, or preparation of the manuscript.

We thank Susanne Rieckmann for excellent technical assistance. We thank Anke Schönherr and Caterina Michetti for organizational matters with SynTKO animals; as well as Christian Hoffmann and Franziska Trnka for assistance in obtaining the necessary animal permits for the SynTKO line. We thank the Electron Microscopy Laboratory of the Institute of Integrative Neuroanatomy and the Core Facility for Electron Microscopy of the Charité for granting us access to their instruments. Finally, we thank Antje Fortströer for careful revision of our manuscript.

