## Supplementary figures and images for "The synapsin-dependent vesicle cluster is crucial for presynaptic plasticity at a glutamatergic synapse in male mice"

### Supplementary Figure 1-1

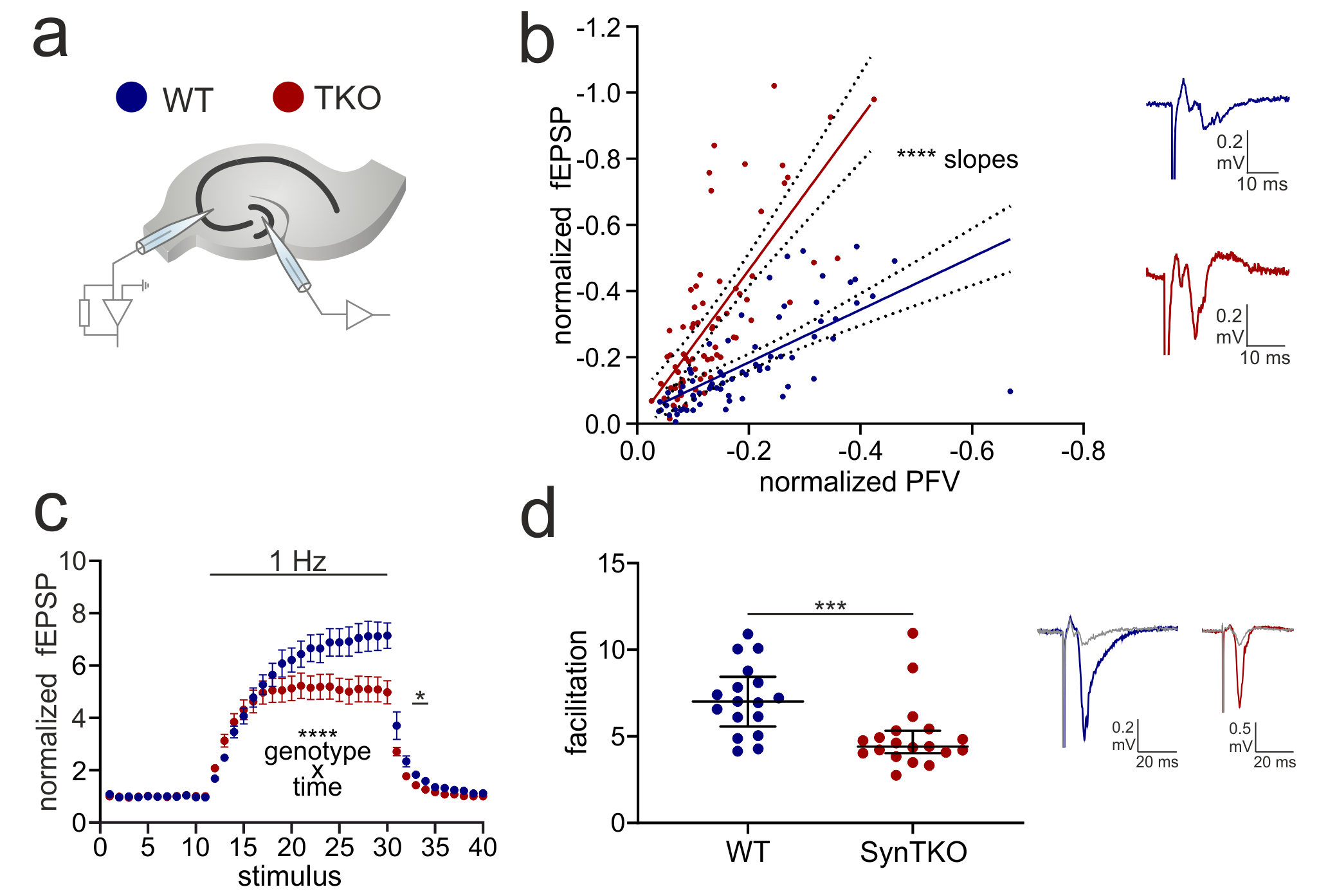

### Supplementary Figure 1-2

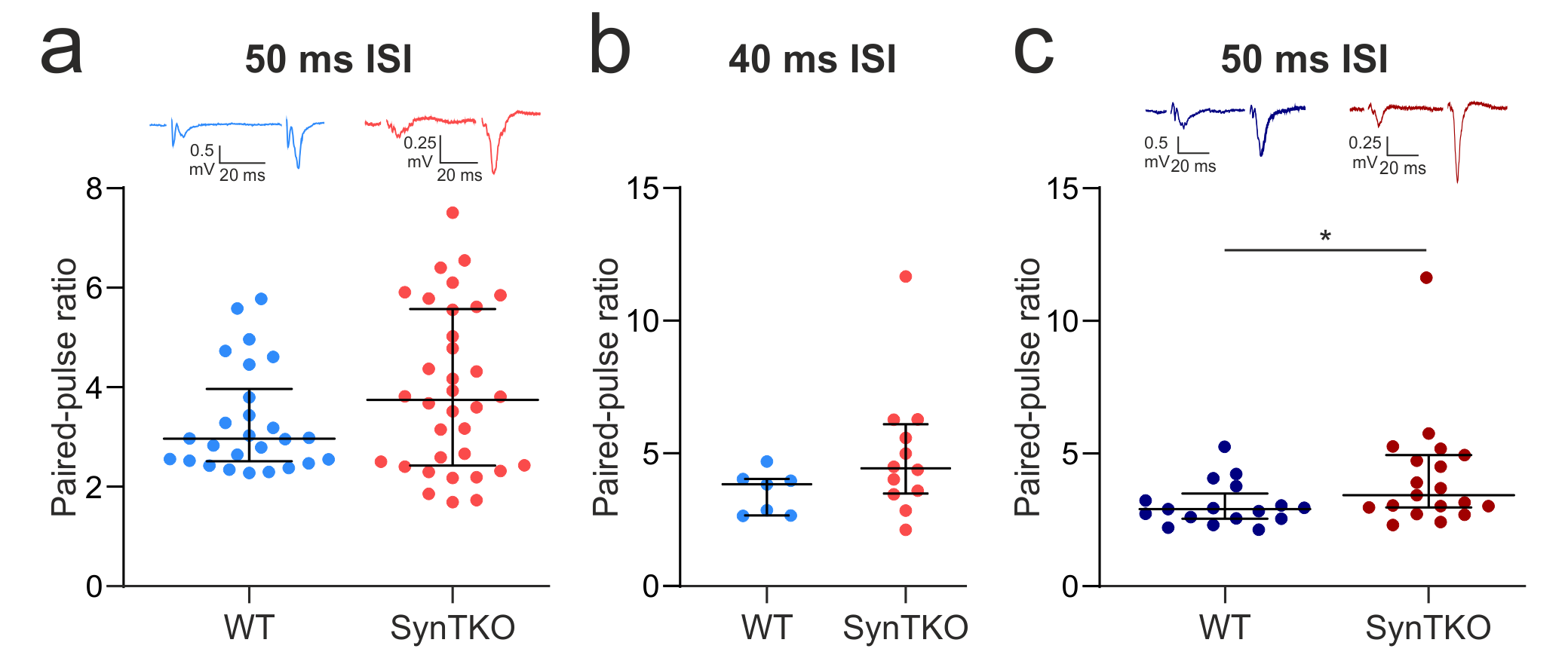

### Supplementary Figure 2-1

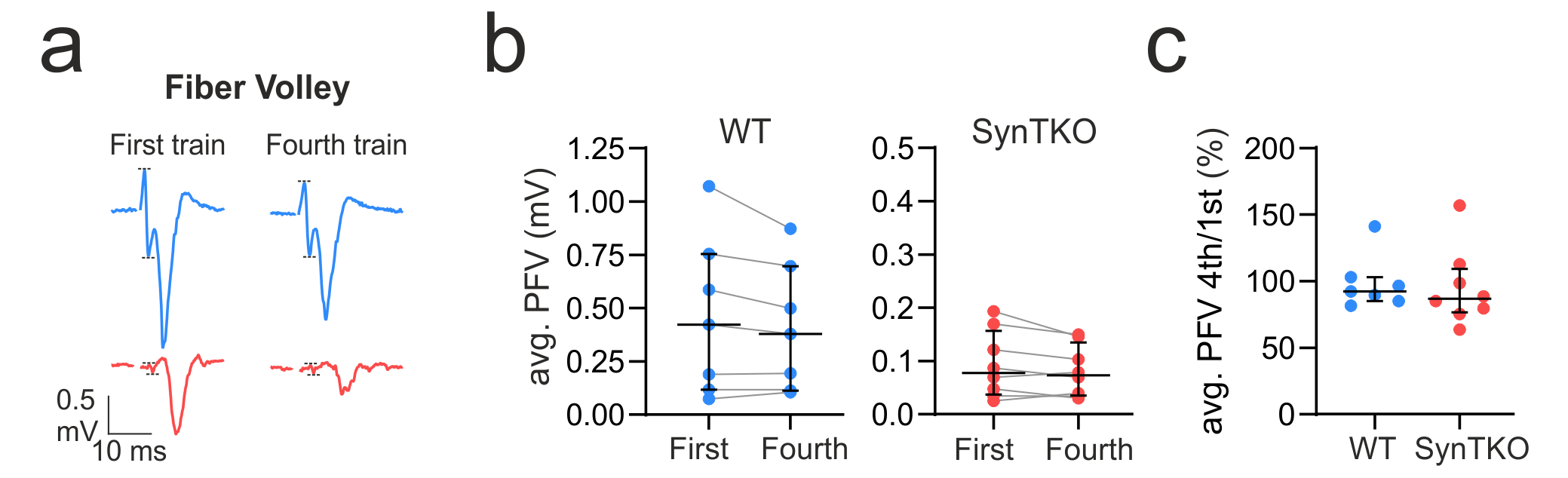
